# The allelic rice immune receptor Pikh confers extended resistance to strains of the blast fungus through a single polymorphism in the effector binding interface

**DOI:** 10.1101/2020.09.05.284240

**Authors:** Juan Carlos De la Concepcion, Josephine H. R. Maidment, Apinya Longya, Gui Xiao, Marina Franceschetti, Mark J. Banfield

## Abstract

Arms race co-evolution drives rapid adaptive changes in pathogens and in the immune systems of their hosts. Plant intracellular NLR immune receptors detect effectors delivered by pathogens to promote susceptibility, activating an immune response that halts colonization. As a consequence, pathogen effectors evolve to escape immune recognition and are highly variable. In turn, NLR receptors are one of the most diverse protein families in plants, and this variability underpins differential recognition of effector variants. The molecular mechanisms underlying natural variation in effector recognition by NLRs are starting to be elucidated. The rice NLR pair Pik-1/Pik-2 recognizes AVR-Pik effectors from the blast fungus *Magnaporthe oryzae*, triggering immune responses that limit rice blast infection. Allelic variation in a heavy metal associated (HMA) domain integrated in the receptor Pik-1 confers differential binding to AVR-Pik variants, determining resistance specificity. Previous mechanistic studies uncovered how a Pik allele, Pikm, has extended recognition to effector variants through a specialized HMA/AVR-Pik binding interface. Here, we reveal the mechanistic basis of extended recognition specificity conferred by another Pik allele, Pikh. A single residue in Pikh-HMA increases binding to AVR-Pik variants, leading to an extended effector response in planta. The crystal structure of Pikh-HMA in complex with an AVR-Pik variant confirmed that Pikh and Pikm use a similar molecular mechanism to extend their pathogen recognition profile. This study shows how different NLR receptor alleles functionally converge to extend recognition specificity to pathogen effectors.

**Author Summary:** Plant pathogens constantly evolve to overcome immune defences and successfully colonize hosts, resulting in some of the most devastating diseases that affect global food production. To defend themselves, plants have evolved a sophisticated immune system that recognizes the presence of different pathogens and triggers immune responses to stop their spread. How plant immune receptors achieve extended recognition to specific pathogen strains and the molecular details of this recognition are just starting to be understood.

In this study, we characterize how an allele of a rice immune receptor achieves a broad-spectrum recognition to effectors from the rice blast fungus. We found that this receptor has evolved a single change that alters the way it binds to different effector variants. This change increases binding affinity to these variants and this is ultimately translated to immune recognition. Interestingly, a different rice immune receptor allele also achieves broad-spectrum effector recognition in a similar way. Therefore, different immune receptor alleles can converge on a similar mechanism to achieve extended recognition to pathogen effectors.

This knowledge has the potential to help to the rational design of plant immune receptors with bespoke resistance to some of the most destructive pathogens. A long-term goal in plant biotechnology.

## Introduction

Plant pathogens cause extensive yield losses in crop harvests worldwide [1]. To ensure successful colonization, pathogens secrete an arsenal of effector molecules that are delivered into host cells to circumvent immune defences and manipulate cell processes, ultimately promoting infection [2]. To counteract these virulence factors, plants have evolved an array of intracellular immune receptors belonging to the nucleotide-binding, leucine-rich repeat (NLR) superfamily that can detect pathogen effectors [3]. Upon recognition, NLRs trigger the activation of immune responses that ultimately lead to localised programmed cell death, stopping the spread of the pathogen [4, 5].

Recognition by immune receptors imposes a strong constraint on pathogens, driving the evolution of new effector variants that escape immune detection. To match this, NLRs are present in large and diverse protein families in plants [6, 7], often with discrete recognition specificity for effector variants [8-10]. As a result, both pathogen effectors and plant NLRs are under strong selection and present signatures of rapid evolution [11-13].

Plant NLRs use diverse mechanisms to recognize pathogen effectors and/or their activities [14, 15]. Multiple plant NLRs harbour non-canonical domains integrated in their architecture. These integrated domains are thought to mimic host proteins targeted by effectors, and serve as baits to mediate pathogen detection [14, 16]. The abundance of integrated domains found across plant genomes suggests that this is an evolutionarily favourable mechanism of pathogen recognition [17-19]. The discovery of integrated domains in plant NLRs facilitates the mechanistic study of effector recognition [20-23] and presents new opportunities to engineer disease resistance [24].

The fungus *Magnaporthe oryzae* causes blast disease in rice [25, 26] and other cereal crops such as barley and wheat [27, 28]. The genome of this pathogen encodes hundreds of putative effectors [29], some of which are recognized by plant NLRs, leading to disease resistance [30]. Paired NLR receptors harbouring integrated domains are particularly prevalent in rice [31, 32], and account for some of the most well-characterised resistance genes against rice blast [33]. The effector complement of different blast strains shows signatures of rapid evolution, including presence/absence polymorphisms [34-37]. This allows the blast pathogen to break genetic resistance, producing disease outbreaks that threaten food production worldwide [1, 27, 38].

AVR-Pik is one of the several rice blast effectors characterized to date [39], and belongs to the <u>M</u>agnaporthe <u>A</u>VRs and To<u>x</u>B like (MAX) effector family, whose members share a similar overall structural scaffold despite divergent sequence [40]. AVR-Pik is recognized in rice by a pair of genetically linked NLRs, Pik-1 and Pik-2 [33, 41]. The sensor NLR Pik-1 harbours an integrated heavy metal associated (HMA) domain that directly binds AVR-Pik triggering immune responses [20, 21]. In contrast to other blast effectors such as AVR1-CO39 and AVR-Pii, which present presence/absence polymorphisms in blast genomes [34, 35, 42], AVR-Pik displays signatures of positive selection, and occurs as multiple effector variants [39, 43]. To date, six AVR-Pik variants (A to F) have been described [39, 44]. Polymorphisms in these variants affect their binding to the Pik-HMA domain and can lead to escape from Pik detection [20, 21, 45].

Pik NLRs exist as an allelic series across rice cultivars. Five different Pik alleles, Pikh, Pikp, Pikm, Piks and Pik* have been described based on their differential response profiles to blast strains [45-49]. The emergence of these Pik alleles has been proposed to follow a linear progression from narrow to broad recognition spectrum (Piks/Pikp → Pik* → Pikm/Pikh), driven by co-evolution with AVR-Pik variants [33, 43, 45]. Interestingly, none of the Pik alleles mediate resistance to blast strains harbouring AVR-PikC or AVR-PikF [44, 45]. Polymorphisms that define the allelic diversity of Pik NLRs are located within the HMA domain, with which the effector interacts [45, 50]. Ultimately, recognition specificity is underpinned by modifications in the HMA binding interface that determine binding to AVR-Pik effectors [20].

The differential recognition of AVR-Pik between Pikp (narrow-spectrum) and Pikm (broad-spectrum) is defined at the structural level via three binding interfaces [20]. Whereas Pikp-HMA interface 2 confers efficient binding and recognition of AVR-PikD, Pikm-HMA interface 3 supports extended effector recognition to AVR-PikD, AVR-PikE and AVR-PikA [20]. Furthermore, incorporating the Pikm-HMA interface 3 into Pikp (Pikp^NK-KE^) extended the binding affinity and recognition profile of Pikp to AVR-PikE and AVR-PikA [24]. Together, these results suggest that Pik alleles have separately evolved distinct molecular mechanisms to ensure efficient effector recognition. Supporting this, HMA domains from allelic Pik receptors cluster into two phylogenetically distinct groups (Bootstrap = 100) that contain either cultivar K60 or Tsuyuake, which harbour Pikp and Pikm resistance, respectively **(Figure 1a, Supplementary Figure 1)** [50]. Each group comprises alleles with narrow and broad recognition specificities to different AVR-Pik variants [45] **(Figure 1b)**, suggesting that diverse Pik alleles have separately evolved broader recognition to AVR-Pik variants.

**Figure 1.**
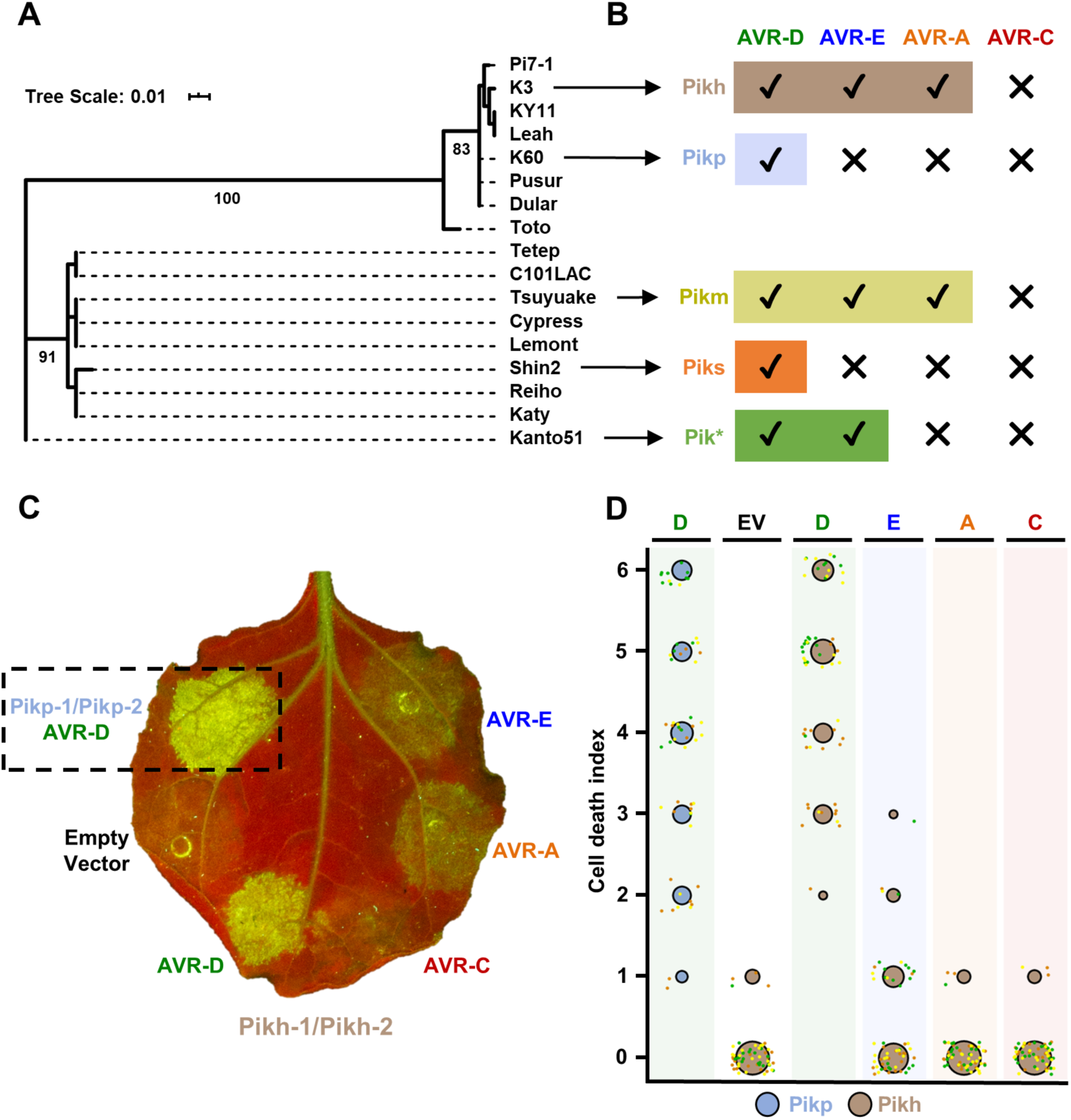
Pikh cell death responses to AVR-Pik variants in *N. benthamiana*. **(A)** Maximum Likelihood Phylogenetic tree of coding sequences of rice Pik-1 HMA domains. The tree was prepared using Interactive Tree Of Life (iTOL) v4 [77]. Cultivar names are placed next to their corresponding branch. Significant bootstrap values (>75) are indicated. **(B)** Schematic representations of immune response profiles of rice cultivars K3 (Pikh), K60 (Pikp), Tsuyuake (Pikm), Shin2 (Piks) and Kanto51 (Pik*) as reported in [45]. **(C)** Representative leaf image showing Pikh-mediated cell death to AVR-Pik variants as autofluorescence under UV light. Pikp-mediated cell death with AVR-PikD is included as a positive control (surrounded by a dashed square), and a spot inoculated with empty vector instead of AVR-Pik effector is included a negative control. **(D)** Cell death assay scoring represented as dot plots. Fluorescence intensity is scored as previously described in [20, 21]. Pikh-mediated cell death is coloured in brown while the Pikp control is coloured in blue. For each sample, all the data points are represented as dots with a distinct colour for each of the three biological replicates; these dots are jittered about the cell death score for visualisation purposes. The size of the centre dot at each cell death value is directly proportional to the number of replicates in the sample with that score. The total number of repeats was 60.

The Pikh allele, present in rice cultivar K3, displays extended recognition of rice blast fungus isolates carrying different AVR-Pik variants [43, 45, 50]. Pikh clusters in the same phylogenetic group as the narrow-spectrum allele Pikp **(Figure 1a, b)**. However, the disease resistance profile of rice cultivar K3 (Pikh) is similar to Tsuyuake (Pikm) [45] **(Figure 1b)**. The only polymorphism between Pikp and Pikh, Asn261Lys, maps to the HMA domain and is contained within binding interface 3 **(Supplementary Figure 1)**. This is the region that underpins extended pathogen recognition in Pikm [20], and is one of the mutations previously shown to extend AVR-Pik recognition profile when introduced in Pikp [24].

Here, we show that the single amino acid polymorphism Asn261Lys in Pikh-HMA increases the binding affinity to AVR-Pik effectors, underpinning the extended recognition of Pikh to AVR-Pik variants. The crystal structure of Pikh-HMA bound to AVR-PikC shows that the Asn261Lys polymorphism in Pikh introduces a Pikm-like interface 3 to aid effector binding. These results demonstrate that Pikh and Pikm have independently converged towards a similar molecular mechanism to confer broad-spectrum resistance to blast strains.

## Results

### Pikh-1/Pikh-2 mediate an extended response to rice blast AVR-Pik effector variants in *N. benthamiana*

*N. benthamiana* is a well-established model system to monitor Pik-mediated cell death in response to AVR-Pik effectors following transient expression via agroinfiltration [20, 21, 24]. We used this system to explore the extended recognition specificity to AVR-Pik effector variants observed for Pikh in rice [45]. For this, we co-expressed Pikh-1 and Pikh-2 (which is 100% identical to Pikp-2) in *N. benthamiana* with either AVR-PikD, AVR-PikE, AVR-PikA or AVR-PikC **(Figure 1c)**. We co-expressed Pikh-1/Pikh-2 with empty vector as a negative control, and Pikp-1/Pikp-2 and AVR-PikD as a positive control **(Figure 1c)**.

In this assay, Pikh shows a robust cell death response to AVR-PikD, a weak response to AVR-PikE, but no response to AVR-PikA or AVR-PikC (comparable to the negative control **(Figure 1c,d)**). The expression of each protein was confirmed by western blot **(Supplementary Figure 2)**. These results show that Pikh has an extended response to AVR-Pik effectors in *N. benthamiana* compared to Pikp, but not to the same extent as previously seen for Pikm [20].

### The Pikh-HMA domain binds to AVR-Pik effectors more strongly than Pikp-HMA

We sought to determine whether the extended recognition mediated by Pikh to strains of *M. oryzae* [43, 45, 50] correlates with an increase in binding of the Pikh-HMA to the AVR-Pik effector variants.

First, we tested the interaction of Pikh-HMA with AVR-Pik variants by yeast-2-hybrid (Y2H), using Pikp-HMA for comparison **(Figure 2a)**. As previously reported, Pikp-HMA interacted with AVR-PikD (depicted by yeast growth and development of blue coloration), while AVR-PikE and AVR-PikA show reduced interaction **(Figure 2a)** [20, 24]. For Pikh-HMA, we observed a similar interaction with AVR-PikE and an increase in the interaction with AVR-PikA, compared to Pikp-HMA **(Figure 2a)**, which was more pronounced after a longer incubation period **(Supplemental figure 3)**. Interestingly, Pikh-HMA displayed interaction with AVR-PikC in this assay **(Figure 2a, Supplemental figure 3)**, but neither Pikp-HMA nor Pikh-HMA displayed any interaction with AVR-PikF. The expression of each protein in yeast was confirmed by western blot **(Supplemental Figure 4)**.

**Figure 2.**
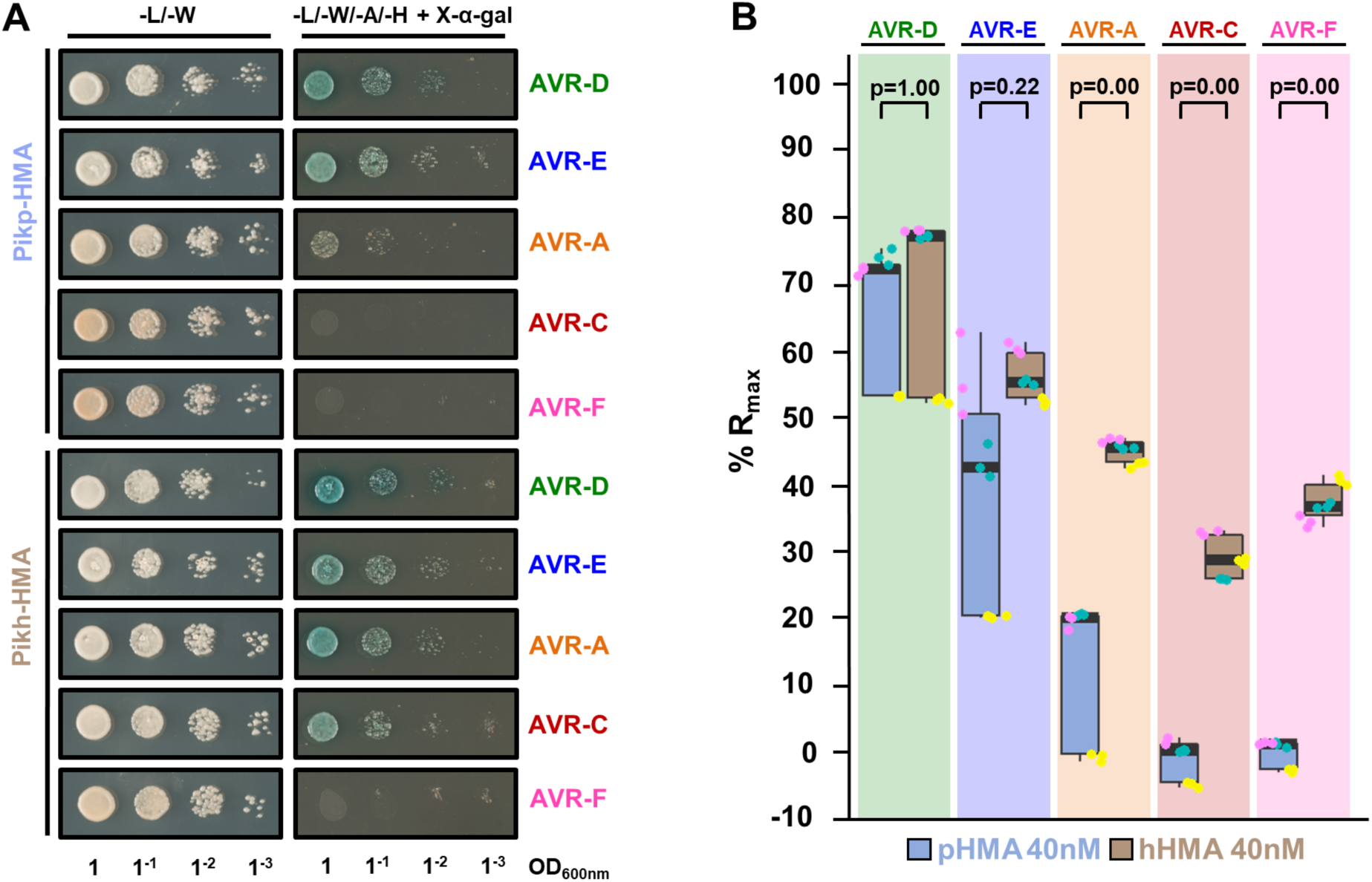
Pikh-HMA has increased binding to AVR-Pik effector alleles in vivo and in vitro. **(A)** Yeast two-hybrid assay of Pikp-HMA and Pikh-HMA with AVR-Pik variants. For each combination of HMA/AVR-Pik, 5μl of yeast were spotted and incubated for ∼60 h in double dropout plate for yeast growth control (left) and quadruple dropout media supplemented with X-α-gal (right). Growth, and development of blue colouration, in the selection plate are both indicative of protein:protein interaction. HMA domains were fused to the GAL4 DNA binding domain, and AVR-Pik alleles to the GAL4 activator domain. Each experiment was repeated a minimum of three times, with similar results. **(B)** Measurement of Pikp-HMA and Pikh-HMA binding to AVR-Pik effector variants by surface plasmon resonance. The binding is expressed as %R_max_ at an HMA concentration of 40 nM. Pikp-HMA and Pikh-HMA are represented by blue and brown boxes, respectively. For each experiment, three biological replicates with three internal repeats each were performed, and the data are presented as box plots. The centre line represents the median, the box limits are the upper and lower quartiles, the whiskers extend to the largest value within Q1 - 1.5× the interquartile range (IQR) and the smallest value within Q3 + 1.5× IQR. All the data points are represented as dots with distinct colours for each biological replicate. “p” is the p-value obtained from statistical analysis and Tukey’s HSD. For results of experiments with 4 and 100 nM HMA protein concentrations, see **Supplemental figure 5**.

**Figure 3.**
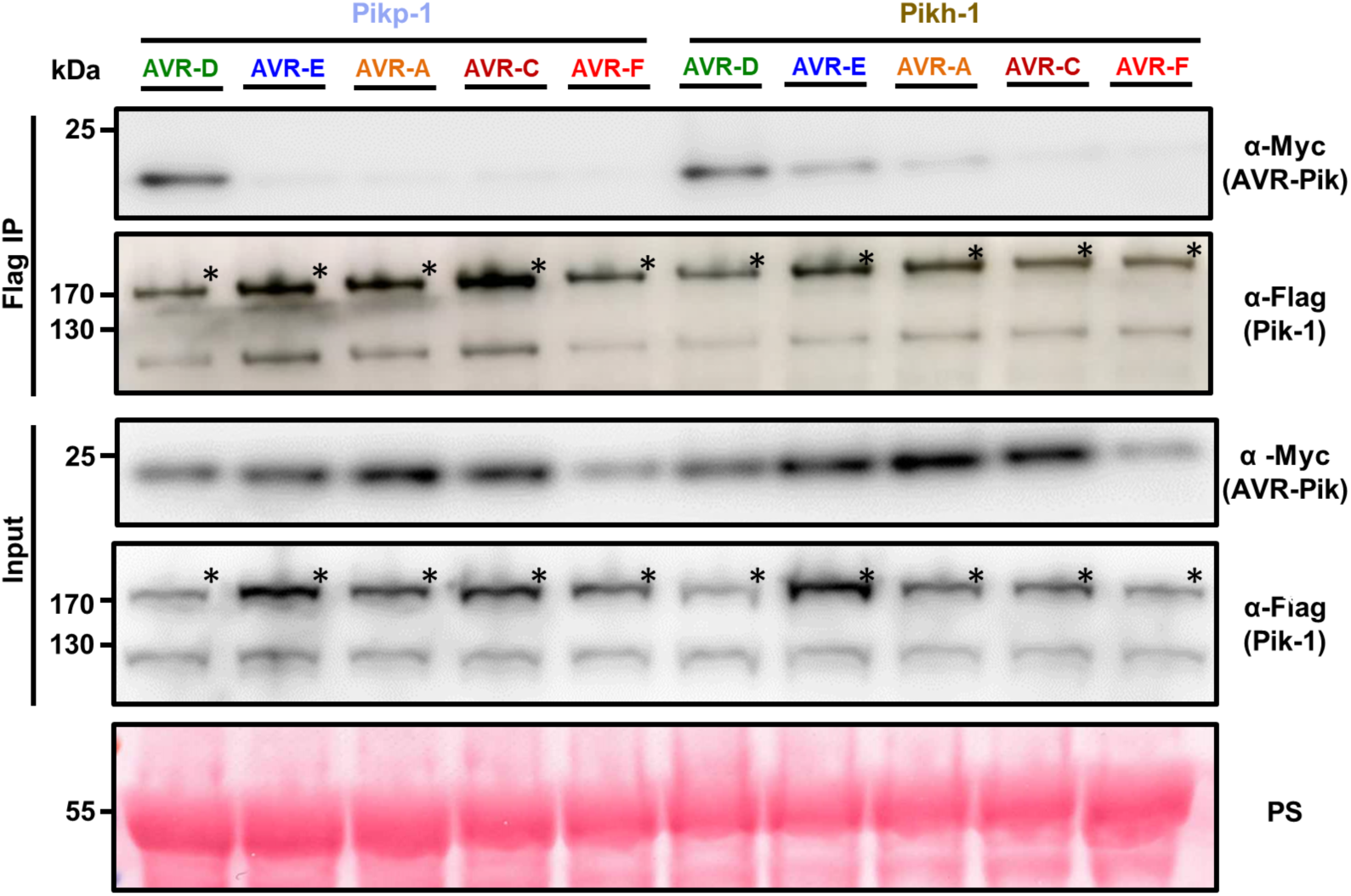
The Asn261Lys polymorphism in Pikh-1 extends association to AVR-PikE and AVR-PikA in planta. Co-immunoprecipitation of full length Pikp-1 and Pikh-1 with AVR-Pik variants. N-terminally 4xMyc tagged AVR-Pik effectors were transiently co-expressed with Pikp-1:6xHis3xFLAG (left) or Pikh-1:6xHis3xFLAG (right) in *N. benthamiana*. Immunoprecipitates (IPs) obtained with M2 anti-FLAG resin and total protein extracts were probed with appropriate antisera. Each experiment was repeated at least three times, with similar results. The asterisks mark the Pik-1 band. Total protein extracts were coloured with Ponceau Staining (PS).

**Figure 4.**
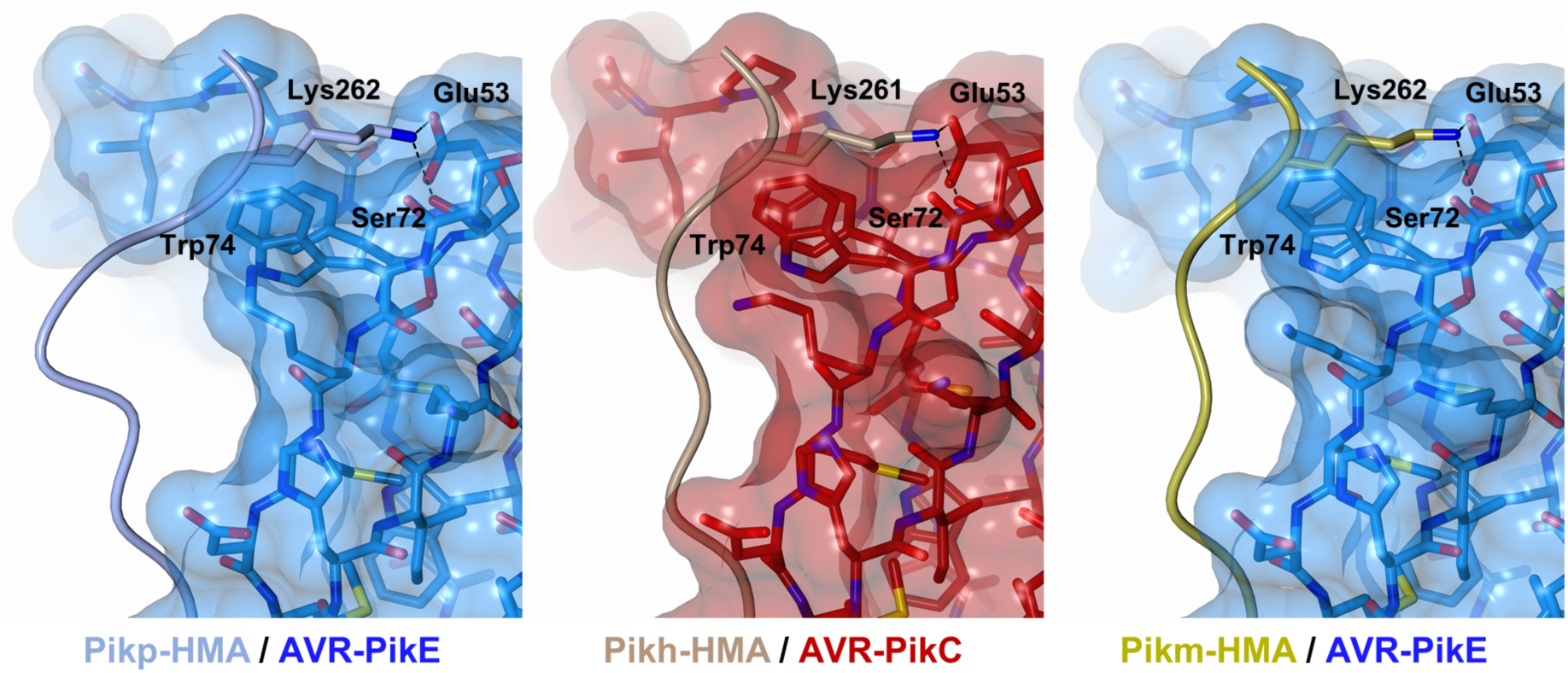
The Pikh-HMA domain adopts a favourable conformation at the effector binding interface. Schematic representation of the conformations adopted by Pikp-HMA (PDB: 6G11), Pikm-HMA (PDB: 6FUB) and Pikh-HMA at interface 3 in complex with AVR-PikE or AVR-PikC. In each panel, the effector is represented in cylinders, with the molecular surface also shown and coloured as labelled. Pik-HMA residues are coloured as labelled and shown as the Cα-worm. For clarity, only the Lys-261/262 side chain is shown. Hydrogen bonds between Lys-261/262 and the effector are represented by dashed black lines. (left) Pikp-HMA bound to AVR-PikE, (middle) Pikh-HMA bound to AVR-PikC, (right) Pikm-HMA bound to AVR-PikE.

Next, we expressed and purified Pikp-HMA and Pikh-HMA domains, and the AVR-Pik variants, in *E. coli* using established protocols as described in the **Materials and Methods** [20, 21, 24]. We used surface plasmon resonance (SPR) to quantitatively measure and compare protein binding [51] **(Figure 2b, Supplemental Figure 5)**. We captured each AVR-Pik variant onto a Biacore NTA chip via a hexahistidine tag at the C-terminus of the effector. Then, we injected either Pikp-HMA or Pikh-HMA at three different concentrations (4 nM, 40 nM and 100 nM), recording the binding level in Response Units (RUs). RUs were then normalised to the theoretical maximum response (R_max_) as described in [51], assuming a 2:1 (Pik-HMA:AVR-Pik) interaction model. This assay showed an increased binding of Pikh-HMA to all the AVR-Pik effectors in vitro, compared with Pikp-HMA **(Figure 2b, Supplemental Figure 5)**. This further confirmed the interaction between Pikh-HMA and AVR-PikC observed in Y2H **(Figure 2a, b)** and, although we did not observe interaction by Y2H, we also detect binding between Pikh-HMA and AVR-PikF by SPR **(Figure 2b, Supplemental Figure 5)**.

**Figure 5.**
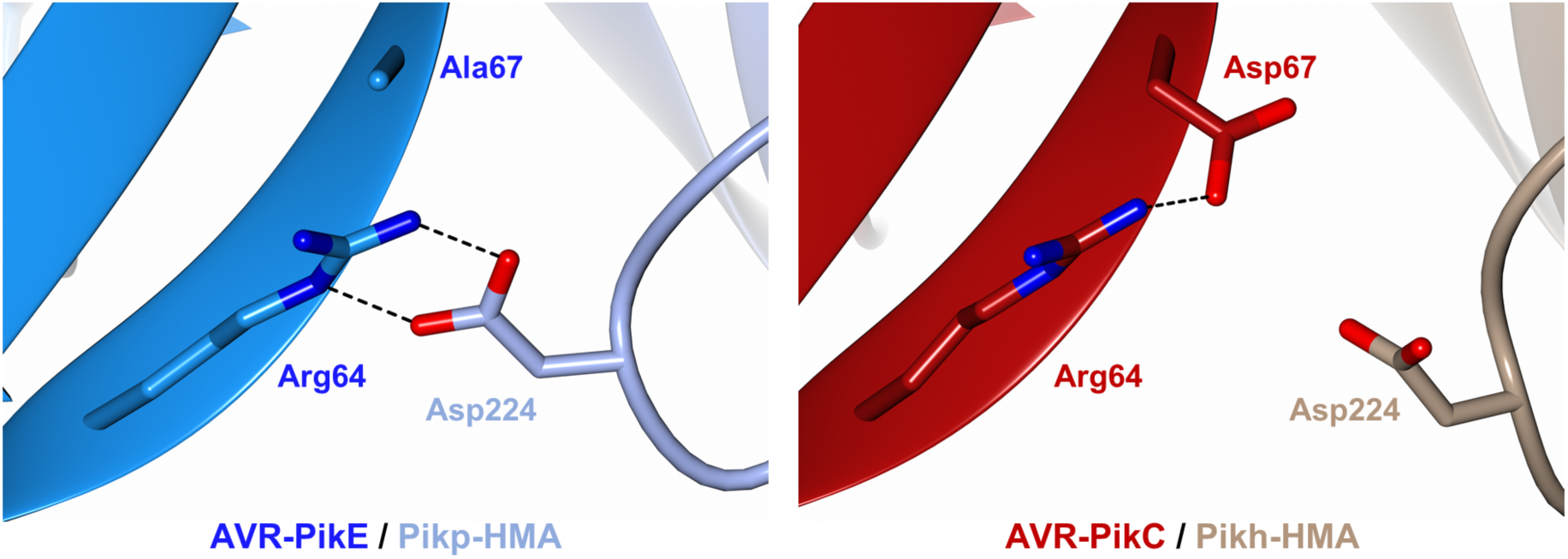
The polymorphic Asp67 in AVR-PikC disrupts hydrogen bonding between the effector and the HMA domain. Close-up views of the position and interactions of Asp224 of the HMA domain in complex with either AVR-PikE (Pikp-HMA (PDB: 6G11), left) or AVR-PikC (Pikh-HMA, right). HMA domains are presented as cartoon ribbons with the side chain of Asp224 displayed as a cylinder; Pikh-HMA and Pikp-HMA are coloured in brown and ice blue, respectively. The effectors are shown in cartoon ribbon representation, with the side chains of Arg64 and Asp67/Ala67 as cylinders. AVR-PikC and AVR-PikE are coloured in crimson and bright blue, respectively. Hydrogen bonds/salt bridges are shown as black dashed lines. For clarity, the N-terminal residues 32 to 52 of the AVR-Pik effector are hidden from the foreground in both structures.

Together, the Y2H and SPR results show that Pikh-HMA displays increased binding to AVR-Pik effector variants compared to Pikp-HMA in vitro. This partially correlates with the extended recognition specificity displayed by the rice cultivar K3 harbouring Pikh [45] **(Figure1b)** and, to a lesser extent, with the cell death assays in *N. benthamiana* **(Figure 1c, d)**. As the only difference between the Pikp and Pikh HMA domains is the single amino acid polymorphism Asn261Lys **(Supplemental Figure 1)**, this amino acid must underpin the increase in effector binding.

### The increased binding of Pikh-HMA to AVR-Pik variants results in an extended effector association in planta

Previous studies have shown that Pik-HMA domain binding to AVR-Pik in yeast and in vitro may not always directly correlate with pathogen recognition by the host, most likely due to lacking the context of the full-length receptor and conditions of the plant cell [20, 24]. Therefore, we investigated the association between full-length Pikh-1 and AVR-Pik effectors in planta. For this, we co-expressed either Pikp-1 or Pikh-1 with each of the AVR-Pik variants in *N. benthamiana*. Pik-1 proteins were subsequently immunoprecipitated and effector association was determined by western blot **(Figure 3)**.

As previously reported, Pikp-1 robustly associates with AVR-PikD, as shown by the signal in the co-IP blot developed with α-Myc tag, but not with AVR-PikE or AVR-PikA **(Figure 3)** [24]. For Pikh-1, we observe a stronger association of full-length Pikh-1 to AVR-PikE and AVR-PikA compared to Pikp-1 **(Figure 3)**. Furthermore, the association levels of AVR-PikD, AVR-PikE and AVR-PikA follow the trend of cell death in *N. benthamiana* **(Figure 1c, d)**. Although Pikh-HMA interacts with AVR-PikC and AVR-PikF in vitro, we found no association between full-length Pikh-1 and either of these effector variants in this assay **(Figure 3)**.

These results suggest that the increased binding of Pikh-HMA to effector variants extends the association of the full-length Pikh-1 receptor to AVR-PikE and AVR-PikA in planta. This correlates with the recognition specificity displayed by rice cultivars harbouring the Pikh allele [45].

### A single polymorphism at the Pikh-HMA effector-binding interface underpins the increased binding to AVR-Pik effectors

To gain a mechanistic understanding of how the Pikh Asn261Lys polymorphism increases the association of the receptor with AVR-Pik effectors, we determined the crystal structure of Pikh-HMA in complex with AVR-PikC.

The complex of Pikh-HMA bound to AVR-PikC was co-expressed and purified from *E. coli* using established protocols [20, 24]. The complex was subsequently crystallised and X-ray diffraction data were collected at the Diamond Light Source (Oxford, UK) to 2.3 Å resolution. Details of protein complex purification, crystallization, data collection, structure solution and model refinement are given in the **Materials and Methods** and **Supplemental Table 1**.

Although this is the first structure of an HMA domain in complex with an effector allele not recognised by any known Pik receptor in rice (AVR-PikC), the overall architecture of the complex is very similar to other HMA/AVR-Pik complexes (RMSD of 0.70 Å when superimposed upon the structure of Pikp-HMA/AVR-PikE, PDB: 6G11) [20, 21, 24] (**Supplemental Figure 6, Supplemental Table 2**).

**Figure 6.**
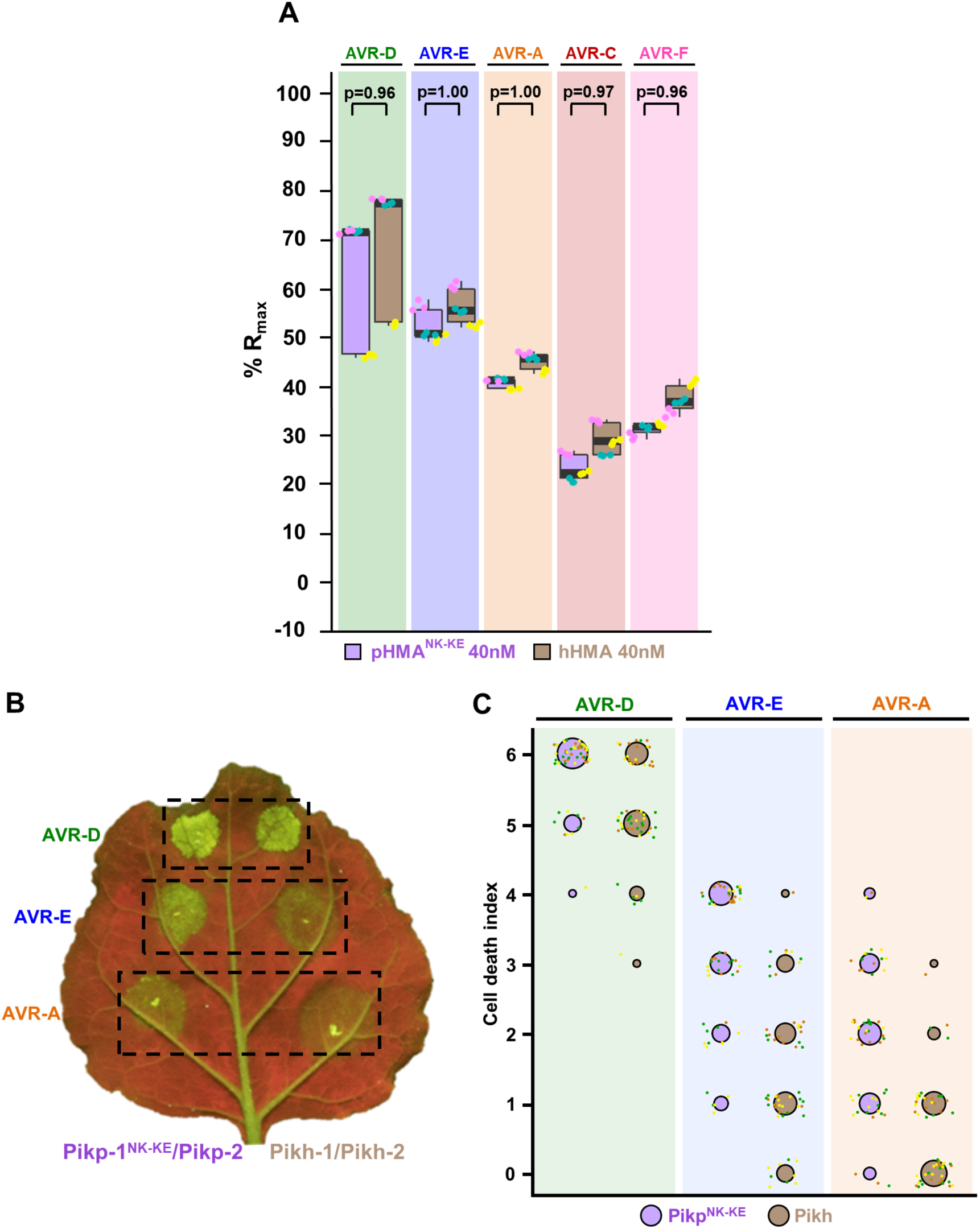
Pikh and Pikp^NK-KE^ display similar binding affinity for AVR-Pik effectors but Pikh shows a reduced cell death response in planta. **(A)** Pikp-HMA^NK-KE^ and Pikh-HMA binding to AVR-Pik effector variants determined by surface plasmon resonance. The binding is expressed as %R_max_ at an HMA concentration of 40 nM. Pikp-HMA^NK-KE^ and Pikh-HMA are represented by purple and brown boxes, respectively. For each experiment, three biological replicates with three internal repeats each were performed and the data are presented as box plots. The centre line represents the median, the box limits are the upper and lower quartiles, the whiskers extend to the largest value within Q1 - 1.5× the interquartile range (IQR) and the smallest value within Q3 + 1.5× IQR. All the data points are represented as dots with distinct colours for each biological replicate. “p” is the p-value obtained from statistical analysis and Tukey’s HSD. Data for Pikh-HMA is also presented in **Figure 2b** and were collected side-by-side at the same time. For results of experiments with 4 and 100 nM HMA protein concentration see **Supplemental figure 8. (B)** Representative leaf image showing a side-by-side cell death assay for Pikp-^NK-KE^ and Pikh with AVR-PikD, AVR-PikE and AVR-PikA. **(C)** Cell death assay scoring represented as dot plots. Fluorescence intensity is scored as previously described in [20, 21]. Cell death mediated by Pikp^NK-KE^ and Pikh is coloured in purple and brown, respectively. For each sample, all the data points are represented as dots with a distinct colour for each of the three biological replicates; these dots are jittered about the cell death score for visualisation purposes. The size of the centre dot at each cell death value is directly proportional to the number of replicates in the sample with that score. The total number of repeats was 57. For statistical analysis of the differences between the cell death mediated by Pikp^NK-KE^ and Pikh see **Supplemental figure 9**.

The polymorphic Asn261Lys is located at the previously described interface 3 [20]. The Pikh-Lys261 residue forms intimate contacts within a pocket formed by AVR-PikC residues Glu53, Tyr71, Ser72 and Trp74 **(Figure 4, middle)**. The position of Lys261 results in a different conformation for the C-terminal region of Pikh-HMA, compared to Pikp-HMA in complex with AVR-PikE **(Figure 4, left)**. However, this conformation is similar to that observed in Pikm-HMA in complex with multiple AVR-Pik effectors. This conformation is thought to extend Pikm recognition to different AVR-Pik variants [20] **(Figure 4, right, Supplemental Figure 7)**.

Altogether, the analysis of the Pikh-HMA/AVR-PikC crystal structure confirms that the single polymorphism Asn261Lys alters the interactions at the HMA/AVR-Pik interface. Further, this data showed that Pikh shares a similar molecular mechanism to extend recognition to AVR-Pik variants with Pikm.

### The polymorphic Asp67 residue in AVR-PikC disrupts hydrogen bonding between AVR-PikC and the HMA domain

To date, there are no reported Pik NLR alleles that confer resistance to rice blast strains carrying AVR-PikC [45]. This effector variant differs from AVR-PikE by a single polymorphism, Ala67Asp, located at the binding interface with the receptor [20, 39]. This polymorphism reduces AVR-PikC binding to Pik-HMA domains and abrogates immune recognition by Pik NLRs [20, 45]. As Pikh-HMA interacts with AVR-PikC in vitro with sufficient affinity to allow co-crystallisation, we were able to investigate the structural basis of how AVR-PikC evades immune recognition.

In the crystal structure of the Pikp-HMA/AVR-PikE complex, the side chain of Asp224 (Pikp-HMA) forms two hydrogen bonds with the side chain of Arg64 (AVR-PikE) **(Figure 5, left)** [20, 24]. By contrast, in the structure of Pikh-HMA/AVR-PikC, the sidechain of Asp67 extends towards the HMA, and the nearby loop containing Asp224 is shifted away from the effector, likely as a consequence of steric clash and/or repulsion due to the matching charges of the two side chains. As a result, there are no hydrogen bonds formed between Asp224 of Pikh-HMA and Arg64 of AVR-PikC **(Figure 5, right)**. Instead, the side chain of Arg64 forms an intramolecular hydrogen bond with the side chain of Asp67. We propose that disruption of hydrogen bonding network at this interface accounts for the lower binding affinity of Pik-HMA domains for AVR-PikC, and the lack of recognition of this effector by Pik NLR proteins in rice.

### Pikh has a similar effector binding and recognition profile to the engineered NLR Pikp^NK-KE^

Based on the structures of Pikm-HMA with AVR-Pik variants, we previously identified a mutant of Pikp, named Pikp^NK-KE^, that extended binding and recognition of this NLR [24]. Interestingly, the Asn261Lys polymorphism found in Pikh is the same as the first position of this double mutant. To better understand the extended recognition phenotype displayed by these NLRs, we compared the Pikh natural variant with the engineered Pikp^NK-KE^.

First, we compared the binding of both Pikh-HMA and Pikp-HMA^NK-KE^ to AVR-Pik variants in vitro **(Figure 6a, Supplemental figure 8)**. We used SPR to quantitatively measure the binding of Pikp-HMA^NK-KE^ to the AVR-Pik variants and compared this with the binding to Pikh-HMA measured above **(Figure 6a, Supplemental Figure 8)**. As previously reported, Pikp-HMA^NK-KE^ showed increased binding to all the AVR-Pik effectors compared with Pikp-HMA, including to AVR-PikC [24]. We also extended this analysis to AVR-PikF and found a similar binding affinity as for AVR-PikC **(Figure 6a, Supplemental Figure 8)**. Overall, the binding levels of Pikp-HMA^NK-KE^ to each of the AVR-Pik effectors were very similar to Pikh-HMA **(Figure 6a, Supplemental Figure 8)**.

We then performed cell death assays in *N. benthamiana* to compare the extent of the immune response of Pikh and Pikp^NK-KE^ to AVR-PikD, AVR-PikE and AVR-PikA **(Figure 6b, c)**. For this, we transiently co-expressed Pikh-1 or Pikp-1^NK-KE^ with Pikp-2 and each of the effectors side-by-side, measuring the cell death response under UV light after 5 days **(Figure 6b, c)**. Pikp^NK-KE^ displayed a clear cell death response to AVR-PikD, AVR-PikE and AVR-PikA with hierarchical levels in the order AVR-PikD > AVR-PikE > AVR-PikA **(Figure 6b, c)** [24], consistent with the binding level of the effectors to the HMA domain **(Figure 6a, Supplemental Figure 8)** [24]. As reported above, Pikh also shows a weak cell death response to AVR-PikE and, in this experiment, to some extent to AVR-PikA **(Figure 1c, d; Figure 6b, c)**. The intensity of Pikh mediated responses in *N. benthamiana* were consistently lower compared to the responses mediated by Pikp^NK-KE^ to each AVR-Pik variant **(Figure 6b, c, Supplemental Figure 9)**. Similar protein accumulation levels were confirmed by western blot **(Supplementary Figure 10)**.

Altogether, these results confirm that the natural polymorphism Asn261Lys in Pikh extends binding and, to some extent, response to AVR-Pik effectors, confirming the results found in the previously characterized Pikp^NK-KE^ mutant, which includes the same mutation [24]. Furthermore, the side-by-side comparison of cell death responses showed that the additional mutation Lys262Glu in the engineered receptor Pikp^NK-KE^ contributes to enhance cell death responses, without affecting the strength of binding to the AVR-Pik effector.

## Discussion

The interplay between pathogen effectors and intracellular immune receptors is one of the most striking examples of arms race co-evolution, and has major biological consequences [11, 52-54]. As their life cycles are commonly far more rapid than the host, pathogens tend to evolve more quickly, producing devastating effects in global agriculture [55]. Therefore, understanding the molecular mechanisms of arms race co-evolution between plants and pathogens has major implications for the development of novel approaches to disease resistance.

Intracellular immune receptors commonly display a narrow recognition specificity to pathogen effectors. Interestingly, NLRs often occur as allelic series with differential effector recognition profiles, which can be governed by direct interaction between the receptor and the pathogen effector. For example, the disease resistance locus Mla encodes allelic NLRs that detect sequence-unrelated effectors from the fungal pathogen *Blumeria graminis f. sp. hordei* (Bgh) [10]. This recognition is mediated by direct interaction [8] which is likely imposing positive selection in the receptor, driving functional diversification [56]. A similar effect can be found in NLRs with integrated domains, as these directly engage with effectors and mediate pathogen recognition. In the case of the Pik locus, the integrated Pik-HMA is the most polymorphic domain [50]. This variation underpins allelic specificity in effector recognition [20] and is likely driven by arms-race co-evolution with the pathogen effector [45]. A linear step-wise model has previously been proposed to illustrate the co-evolutionary dynamics between AVR-Pik effectors and Pik resistance alleles [33, 43, 45]. However, interactions between the allelic AVR-Pik/Pik interactions are more complex, possibly involving differential co-evolution between allelic receptors and their cognate effector variants [20].

Two AVR-Pik variants, AVR-PikC and AVR-PikF, evade recognition by all Pik alleles characterized to date. The polymorphisms defining each of these effectors indicates that they have separately emerged from AVR-PikE and AVR-PikA, respectively [39, 44, 45]. Therefore, there are at least two branches in the evolution of AVR-Pik effectors towards evasion of Pik-mediated immunity. Similarly, the Pik NLR alleles fall in two phylogenetically distinct groups based on their HMA domains. As each group contains members displaying narrow- and broad-spectrum recognition of AVR-Pik alleles, it is likely that extended AVR-Pik recognition phenotypes have evolved separately. This is consistent with previous studies showing that Pikp and Pikm-HMA domains use different interfaces to efficiently bind AVR-Pik effectors [20].

However, it is intriguing to note that Pikm and Pikh appear to have convergently evolved towards the same molecular mechanism to extend effector recognition specificity, by having a lysine residue one position towards the N-terminus in their protein sequence, compared with the narrow-spectrum allele Pikp. This was also the outcome of the Pikp-1^NK-KE^ mutation to artificially extend the recognition spectrum of Pikp through structure-guided engineering [24]. Having a shared polymorphism in a natural (Pikh) and engineered (Pikp-1^NK-KE^) NLR towards expanded effector recognition is perhaps not surprising. This exemplifies how protein engineering approaches can be informed by natural variation in NLR immune receptors, and highlights how polymorphisms that enhance disease resistance can be found in the germplasm of both elite crop varieties and wild relatives [57, 58]. Indeed, mining and characterization of the allelic diversity of integrated domains has the potential to reveal new sources of resistance.

Comparison of the Pikh-HMA and Pikp^NK-KE^-HMA binding to AVR-Pik effectors in vitro and Pikh- and Pikp^NK-KE^-mediated cell death responses in vivo shows that the engineered variant has the potential to perform better in conferring disease resistance in rice as it displays enhanced response in *N. benthamiana*. To date though, it is unknown whether Pikp^NK-KE^ confers a resistance profile to blast strains in rice similar to Pikh [45]. However, these data support the hypothesis that engineering integrated domains of NLR proteins can be used to deliver resistance in crops. But why does the presence of a Glu residue (Pikp^NK-KE^), rather than a Lys residue (Pikh), at position 262 enhance Pik NLR activity? Previous structural studies showed that the side chain of Glu262 is directed away from the binding interface with the effector [24], and would not be expected to influence interaction directly. This is supported by results from SPR experiments presented here **(Figure 6a, Supplemental Figure 8)**. Therefore, Lys262Glu may affect the activation of Pikp-1^NK-KE^ through a mechanism downstream of effector binding. The CC-NLR ZAR1 has recently been shown to oligomerize and form resistosomes upon activation [59]. Given the presence of a MADA motif in the CC domains of ZAR1 and the helper NLR Pik-2, we hypothesise that Pik-2 may use a similar mechanism to trigger cell death in plant cells [60]. Thus, the integrated HMA domain of Pik-1 may make intra and/or intermolecular interactions with other domains in the sensor or helper NLR. Indeed, the integrated WRKY domain present in the Arabidopsis NLR RRS1 has been shown to regulate NLR activation through association with other domains of the NLR [61]. This could explain how polymorphisms that do not alter the strength of effector binding can influence the outcome of immune responses. However, little is known about the intra- and intermolecular interactions that translate effector binding into activation of cell death in the Pik NLR pair.

Polymorphisms in effectors that evade detection by plant immune systems provide a selective advantage to the pathogen. To date, there are two alleles of AVR-Pik that are not recognised in rice by any naturally occurring Pik variant, AVR-PikC and AVR-PikF [44, 45]. AVR-PikC differs from AVR-PikE by a single polymorphism, Ala67Asp. Despite the fact that AVR-PikC is not recognised by Pikh in planta, we were still able to form a Pikh-HMA/AVR-PikC complex in vitro and obtain its crystal structure. This revealed that the Ala67Asp change disrupts an intermolecular hydrogen bonding network, likely showing how AVR-PikC escapes recognition by Pik. This information will inform future engineering efforts to develop Pik receptors that confer disease resistance to blast isolates containing currently unrecognized effector alleles. For example, such efforts could use Pikp^NK-KE^ or Pikh as a scaffold, adding additional mutations that further enhance binding to AVR-Pik effectors and lead to broader resistance to blast disease in rice.

## Accession codes

Structure of Pikh-HMA/AVR-PikC, and the data used to derive this, have been deposited at the Protein DataBank (PDB) with accession code 7A8X.

### Acknowledgements

This work was supported by the UKRI Biotechnology and Biological Sciences Research Council (BBSRC) Norwich Research Park Biosciences Doctoral Training Partnership, UK [grant BB/M011216/1]; the UKRI BBSRC, UK [grants BB/P012574, BB/M02198X]; the European Research Council [ERC; proposal 743165]; the John Innes Foundation; The Thailand Research Fund through The Royal Golden Jubilee Ph.D. Program [PHD/0152/2556]. We would like to thank Andrew Davies and Phil Robinson from JIC Scientific Photography for the UV pictures of the cell death assays, Professor Dan MacLean from the Sainsbury Laboratory (Norwich, UK) Bioinformatics Team for advice on statistical analysis, Dr. Clare Stevenson and Professor David Lawson from the JIC crystallography platform for technical support in protein crystallization and X-ray data collection. We also thank Professor Sophien Kamoun for discussions.

## Materials and Methods

### Gene cloning

For in vitro studies, Pikh-HMA (encompassing residues 186 to 263) was generated by introducing the Asn262Lys mutation in Pikp-HMA by site-directed mutagenesis, followed by cloning into pOPIN-M [62]. Wild-type Pikp-HMA, Pikp-HMA^NK-KE^, and AVR-Pik expression constructs used in this study are as described in [20, 24].

For Y2H, we cloned Pikh-HMA into pGBKT7 using In-Fusion cloning (Takara Bio USA), following the manufacturer’s protocol. Wild-type Pikp-HMA domain in pGBKT7 and AVR-Pik effector variants in pGADT7 used were generated as described in [20]. The *M. oryzae* effector AVR-PikF was cloned into pGADT7 using In-fusion cloning as described above.

For protein expression in planta, the Pikh-HMA domain was generated by introducing the mutation in a reverse primer for PCR. This domain was then assembled into a full-length NLR construct using Golden Gate cloning [63] and into the plasmid pICH47742 with a C-terminal 6xHis/3xFLAG tag. Expression was driven by the *A. tumefaciens* Mas promoter and terminator. Full-length Pikp-1, Pikp-2, and AVR-Pik variants used were generated as described in [20, 44].

All DNA constructs were verified by sequencing.

### Expression and purification of proteins for in vitro binding studies

6xHis-MBP-tagged Pikp-HMA, Pikh-HMA and Pikp-HMA^NK-KE^ were produced in *E. coli* SHuffle cells [64] using the protocol previously described in [20, 24]. Cell cultures were grown in auto induction media [65] at 30°C for 5 – 7hrs and then at 16°C overnight. Cells were harvested by centrifugation and re-suspended in 50 mM HEPES pH 7.5, 500 mM NaCl, 50 mM glycine, 5% (vol/vol) glycerol, 20 mM imidazole supplemented with EDTA-free protease inhibitor tablets (Roche). Cells were sonicated and, following centrifugation at 40000 x *g* for 30 min, the clarified lysate was applied to a Ni^2+^-NTA column connected to an AKTA Xpress purification system (GE Healthcare). Proteins were step-eluted with elution buffer (50 mM HEPES pH 7.5, 500 mM NaCl, 50 mM Glycine, 5% (vol/vol) glycerol, 500 mM imidazole) and directly injected onto a Superdex 75 26/600 gel filtration column pre-equilibrated 20mM HEPES pH 7.5, 150 mM NaCl. Purification tags were then removed by incubation with 3C protease (10 μg/mg fusion protein) overnight at 4°C followed by passing through tandem Ni^2+^-NTA and MBP Trap HP columns (GE Healthcare). The flow-through was concentrated as appropriate and loaded on a Superdex 75 26/600 gel filtration column for final purification and buffer exchange into 20 mM HEPES pH 7.5, 150 mM NaCl.

AVR-Pik effectors, with a 3C protease-cleavable N-terminal SUMO tag and a non-cleavable C-terminal 6xHis tag, were produced in and purified from *E. coli* SHuffle cells as previously described [20, 21, 24]. All protein concentrations were determined using a Direct Detect® Infrared Spectrometer (Merck).

### Co-expression and purification of Pikh-HMA and AVR-PikC for crystallisation

Pikh-HMA was co-expressed with AVR-PikC in *E. coli* SHuffle cells following co-transformation of pOPIN-M:Pikh-HMA and pOPIN-A:AVR-PikC (which were prepared as described in [20, 24]). Cells were grown in autoinduction media (supplemented with both carbenicillin and kanamycin), harvested, and processed as described as above. Protein concentrations were measured using a Direct Detect® Infrared Spectrometer (Merck).

### Crystallization, data collection and structure solution

For crystallization, Pikh-HMA in complex with AVR-PikC was concentrated to ∼10 mg/ml following gel filtration. Sitting drop vapor diffusion crystallization trials were set up in 96 well plates, using an Oryx nano robot (Douglas Instruments, United Kingdom). Plates were incubated at 20°C, and crystals typically appeared after 24 - 48 hours. For data collection, all crystals were harvested from the Morpheus® HT-96 screen (Molecular Dimensions), and snap-frozen in liquid nitrogen. Crystals used for data collection appeared in Morpheus® HT-96 condition F1 [0.12 M Monosaccharides (0.2M D-Glucose; 0.2M D-Mannose; 0.2M D-Galactose; 0.2M L-Fucose; 0.2M D-Xylose; 0.2M N-Acetyl-D-Glucosamine); 0.1 M Buffer system 1 (1 M Imidazole; MES monohydrate (acid)) pH 6.5; 50% v/v Precipitant mix 1 (40% v/v PEG 500; MME; 20 % w/v PEG 20000)].

X-ray data sets were collected at the Diamond Light Source using beamline i04 (Oxford, UK). The data were processed using the autoPROC pipeline [66] as implemented in CCP4i2 [67]. The structures were solved by molecular replacement with PHASER [68] using the coordinates of AVR-PikC and a dimer of Pikp-HMA^NK-KE^ (PDB: 7A8W) as the model. The final structures were obtained through iterative cycles of manual rebuilding and refinement using COOT [69] and REFMAC5 [70], as implemented in CCP4i2 [67]. Structures were validated using the tools provided in COOT and MOLPROBITY [71].

### Protein-protein interaction: Yeast-2-hybrid analyses

To detect protein–protein interactions between Pikh-HMA and AVR-Pik effectors by Yeast Two-Hybrid, we used the Matchmaker® Gold System (Takara Bio USA). We generated a plasmid encoding Pikh-HMA in pGBKT7 and co-transformed it into chemically competent Y2HGold cells (Takara Bio, USA) with the individual AVR-Pik variants in pGADT7 as described previously [20, 24]. Single colonies grown on selection plates were inoculated in 5 ml of SD^-Leu-Trp^ and grown overnight at 30°C. Saturated culture was then used to make serial dilutions of OD_600_ 1, 10^−1^, 10^−2^, 10^−3^, respectively. 5 μl of each dilution was then spotted on a SD^-Leu-Trp^ plate as a growth control, and on a SD^-Leu-Trp-Ade-His^ plate containing X-α-gal (Takara Bio, USA). Plates were imaged after incubation for 60 - 72 hr at 30°C unless otherwise stated. Each experiment was repeated a minimum of 3 times, with similar results.

To confirm protein expression in yeast, total protein extracts from transformed colonies were produced by incubating the cells at 95°C for 10 minutes in LDS Runblue® sample buffer. Samples were centrifuged and the supernatant was subjected to SDS-PAGE gels prior to western blotting. The membranes were probed with anti-GAL4 DNA-BD (Sigma) for the HMA domains in pGBKT7 and anti-GAL4 activation domain (Sigma) antibodies for the AVR-Pik effectors in pGADT7.

### Protein-protein interaction: Surface plasmon resonance

A detailed protocol of the surface plasmon resonance (SPR) experiments can be found in [51]. In brief, experiments were performed on a Biacore T200 system (GE Healthcare) using an NTA sensor chip (GE Healthcare). The system was maintained at 25°C, and a flow rate of 30 μl/min was used. All proteins were prepared in SPR running buffer (20 mM HEPES pH 7.5, 860 mM NaCl, 0.1% Tween 20). C-terminally 6xHis-tagged AVR-Pik variants were immobilised on the chip, giving a response of 250 ± 50 RU. The sensor chip was regenerated between each cycle with an injection of 30 μl of 350 mM EDTA.

For all the assays, the level of binding was expressed as a percentage of the theoretical maximum response (R_max_) normalized to the amount of ligand immobilized on the chip. The cycling conditions were the same as used in [20, 24]. For each measurement, in addition to subtracting the response in the reference cell, a further buffer-only subtraction was made to correct for bulk refractive index changes or machine effects [72]. SPR data was exported and plotted using R v3.4.3 (https://www.r-project.org/) and the function ggplot2 [73]. Each experiment was repeated 3 times, with each replicate including 3 internal repeats.

### Protein-protein interaction: In planta co-immunoprecipitation (co-IP)

Transient gene expression in planta for co-IP was performed by delivering T-DNA constructs with *Agrobacterium tumefaciens* GV3101 strain (C58 (rifR) Ti pMP90 (pTiC58DT-DNA) (gentR) Nopaline(pSoup-tetR)) into 4-week old *N. benthamiana* plants grown at 22–25°C with high light intensity. *A. tumefaciens* strains carrying Pikp-1 or Pikh-1 were mixed with strains carrying the corresponding AVR-Pik effectors, at OD_600_ 0.2 each, in agroinfiltration medium (10 mM MgCl_2_, 10 mM 2-(N-morpholine)-ethanesulfonic acid (MES), pH 5.6), supplemented with 150 μM acetosyringone. For detection of complexes in planta, leaf tissue was collected 3 days post infiltration (dpi), frozen, and ground to fine powder in liquid nitrogen using a pestle and mortar. Leaf powder was mixed with 2x weight/volume ice-cold extraction buffer (10% glycerol, 25 mM Tris pH 7.5, 1 mM EDTA, 150 mM NaCl, 2% w/v PVPP, 10 mM DTT, 1x protease inhibitor cocktail (Sigma), 0.1% Tween 20 (Sigma)), centrifuged at 4,200 x *g*/4 °C for 30 min, and the supernatant was passed through a 0.45μm Minisart® syringe filter. The presence of each protein in the input was determined by SDS-PAGE/western blot. Pik-1 and AVR-Pik effectors were detected probing the membrane with anti-FLAG M2 antibody (SIGMA) and anti-c-Myc monoclonal antibody (Santa Cruz), respectively. For immunoprecipitation, 1.5 ml of filtered plant extract was incubated with 30 μl of M2 anti-FLAG resin (Sigma) in a rotatory mixer at 4°C. After three hours, the resin was pelleted (800 x *g*, 1 min) and the supernatant removed. The pellet was washed and resuspended in 1 ml of IP buffer (10% glycerol, 25 mM Tris pH 7.5, 1 mM EDTA, 150 mM NaCl, 0.1% Tween® 20 (Sigma)) and pelleted again by centrifugation as before. Washing steps were repeated 5 times. Finally, 30 μl of LDS Runblue® sample buffer was added to the agarose and incubated for 10 min at 70°C. The resin was pelleted again, and the supernatant loaded on SDS-PAGE gels prior to western blotting. Membranes were probed with anti-FLAG M2 (Sigma) and anti c-Myc (Santa Cruz) monoclonal antibodies. Each experiment was repeated at least 3 times.

### *N. benthamiana* cell death assays

*A. tumefaciens* GV3101 (C58 (rifR) Ti pMP90 (pTiC58DT-DNA) (gentR) Nopaline(pSoup-tetR)) carrying Pikp-1, Pikh-1 or Pikp-1^NK-KE^ were resuspended in agroinfiltration media (10 mM MES pH 5.6, 10 mM MgCl_2_ and 150 μM acetosyringone) and mixed with *A. tumefaciens* GV3101 carrying Pikp-2, AVR-Pik effectors, and P19 at OD_600_ 0.4, 0.4, 0.6 and 0.1, respectively. 4-weeks old *N. benthamiana* leaves were infiltrated using a needleless syringe. Leaves were collected at 5 dpi and photographed under visible and UV light.

### Cell death scoring: UV autofluorescence

Detached leaves were imaged at 5 dpi from the abaxial side of the leaves for UV fluorescence images. Photos were taken using a Nikon D4 camera with a 60mm macro lens, ISO set 1600 and exposure ∼10secs at F14. The filter is a Kodak Wratten No.8 and white balance is set to 6250 degrees Kelvin. Blak-Ray® longwave (365nm) B-100AP spot light lamps are moved around the subject during the exposure to give an even illumination. Images shown are representative of three independent experiments, with internal repeats. The cell death index used for scoring is as presented previously [21]. Dotplots were generated using R v3.4.3 (https://www.r-project.org/) and the graphic package ggplot2 [73]. The size of the centre dot at each cell death value is directly proportional to the number of replicates in the sample with that score. All individual data points are represented as dots.

### Phylogenetic analysis

Multiple sequence alignment of the coding sequences of 17 Pik-HMA domains obtained from [50] was performed in Clustal Omega [74]. The phylogenetic tree for the Pik-HMA coding sequences was calculated using the Maximum likelihood method and Tamura-Nei model [75] in MEGA X [76]. The tree with the highest log likelihood (−596.61) is shown. Initial trees for the heuristic search were obtained automatically by applying Neighbour-Joining and BioNJ algorithms to a matrix of pairwise distances estimated using the Maximum Composite Likelihood (MCL) approach, and then selecting the topology with superior log likelihood value. A discrete Gamma distribution was used to model evolutionary rate differences among sites (5 categories +G, parameter = 0.4524)). Codon positions included were 1^st^+2^nd^+3^rd^+Noncoding. There were a total of 246 positions in the final dataset. The tree was represented using Interactive Tree Of Life (iTOL) v4 [77].

### Statistical analyses

Qualitative cell-death scoring from autofluorescence was analysed using estimation methods [78] and visualized with estimation graphics using the besthr R library [79]. All cell-death scores in samples under comparison were ranked, irrespective of sample. The mean ranks of the control and test sample were taken and a bootstrap process was begun on ranked test data, in which samples of equal size to the experiment were replaced and the mean rank calculated. After 1000 bootstrap samples, rank means were calculated, a distribution of the mean ranks was drawn and its 2.5 and 97.5 quantiles calculated. If the mean of the control data is outside of these boundaries, the control and test means were considered to be different.

Quantitative R_max_ data from SPR assays were analysed by preparing a linear mixed effects model of sample on SPR. Post-hoc comparisons were performed for sample contrasts using Tukey’s HSD method in the R package nlme [80] and in lsmeans [81].

## Figures

**Supplemental figure 1.**
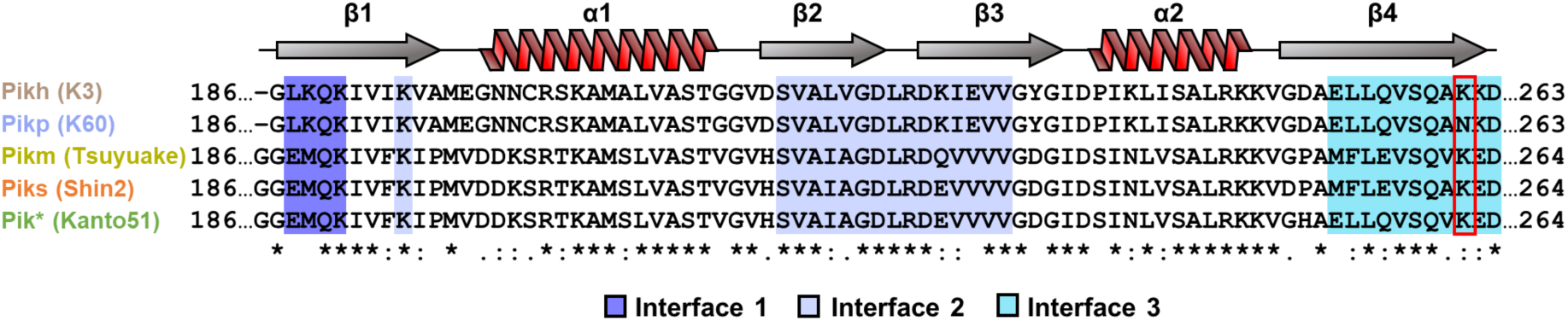
Amino acid sequence alignment of HMA domains of rice cultivars harbouring different Pik alleles. Amino acid sequence alignment of Pikh-1, Pikp-1, Pikm-1, Piks-1 and Pik*-1. Secondary structure features of the HMA fold are shown above, and the residues located to the binding interfaces as described in [20] are highlighted. The Pikh-HMA polymorphic position (residue 261), located in binding interface three, is indicated in a red square.

**Supplemental figure 2.**
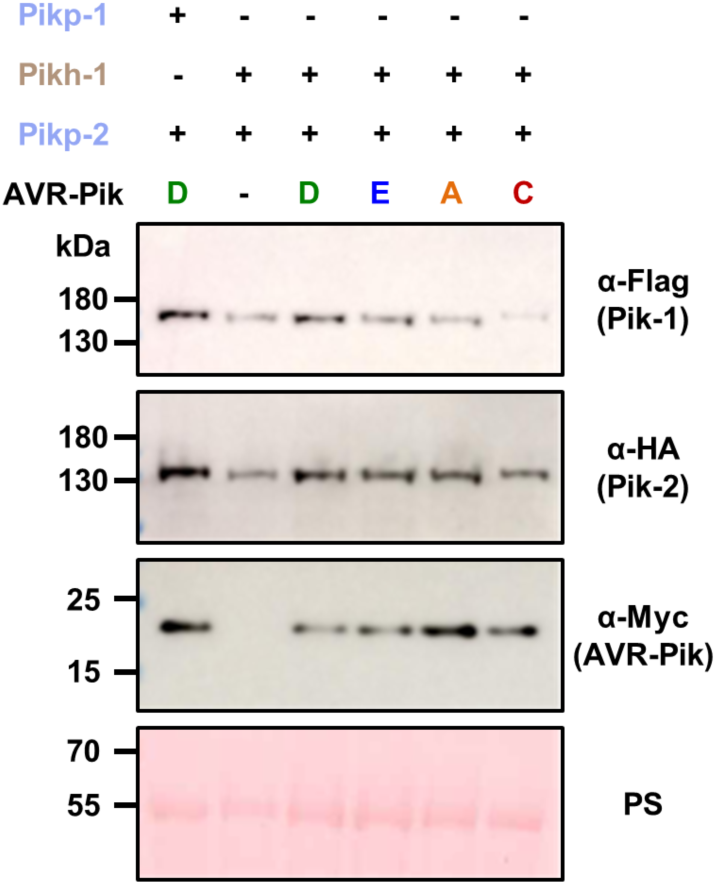
Western blots confirming the accumulation of proteins in *N. benthamiana* cell death assays. Plant lysate was probed for the expression of Pikh-1, Pikp-2 (100% identical to Pikh-2) and AVR-Pik effectors using anti-FLAG, anti-HA and anti-Myc antisera, respectively. Accumulation of the control Pikp-1/Pikp-2/AVR-PikD proteins were also measured as a comparison. Total protein extracts were visualized by Ponceau Staining (PS).

**Supplemental figure 3.**
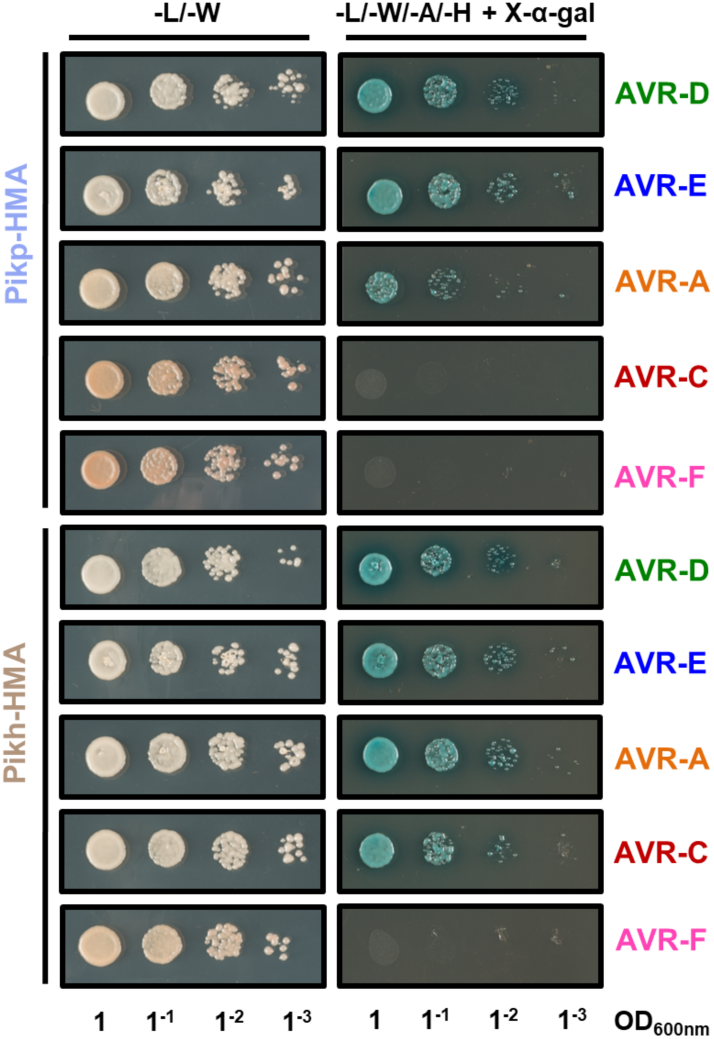
Yeast two-hybrid assay of Pikp-HMA and Pikh-HMA with AVR-Pik variants following an extended incubation. For each combination of HMA/AVR-Pik, 5μl of yeast were spotted and incubated for ∼84 h in double dropout plate for yeast growth control (left) and quadruple dropout media supplemented with X-α-gal (right). Growth, and development of blue colouration, in the selection plate are both indicative of protein:protein interaction. HMA domains were fused to the GAL4 DNA binding domain, and AVR-Pik alleles to the GAL4 activator domain. Each experiment was repeated a minimum of three times, with similar results.

**Supplemental figure 4.**
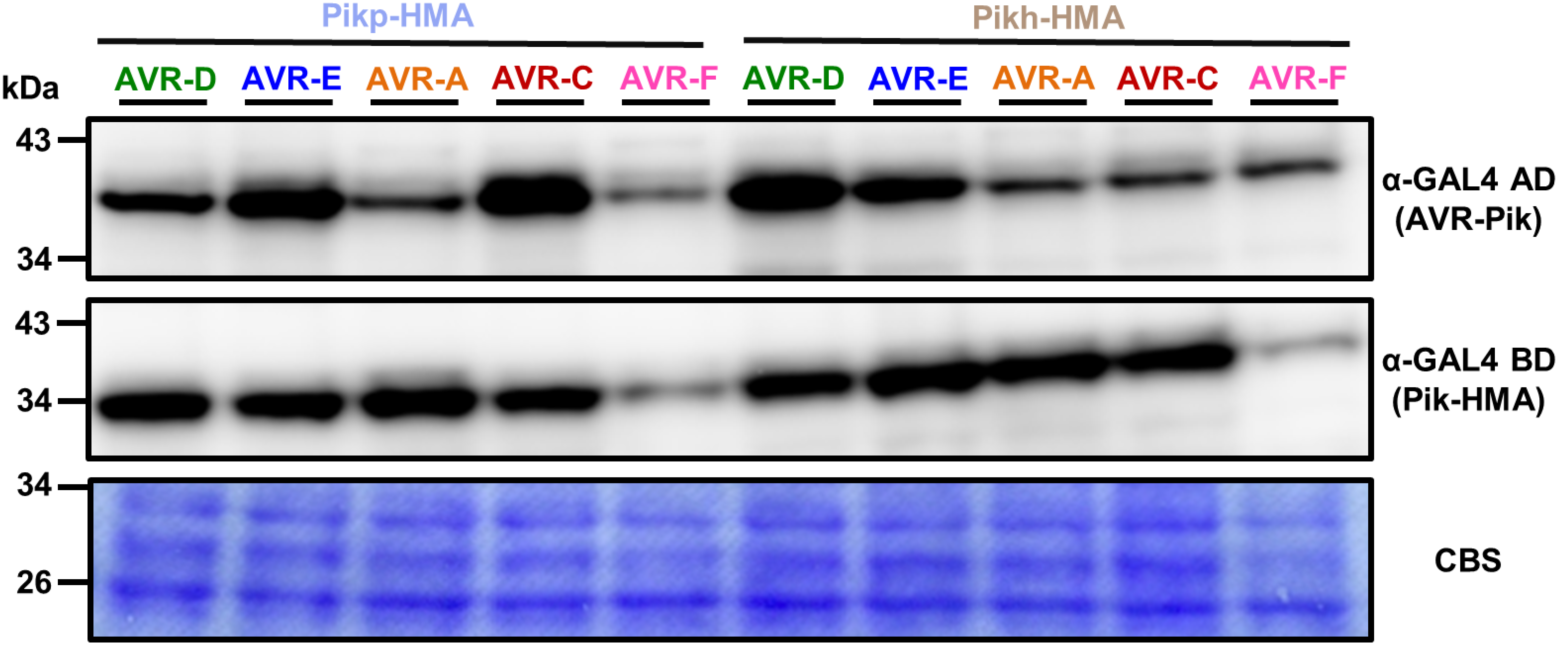
Accumulation of proteins in yeast-two-hybrid assays analysed by Western blot. Yeast lysate was probed for the expression of AVR-Pik effectors and HMA domains using anti-GAL4 activation domain (AD) and anti-GAL4 DNA binding domain (BD) antibodies, respectively. Total protein extracts were coloured with Coomassie Blue Stain (CBS).

**Supplemental figure 5.**
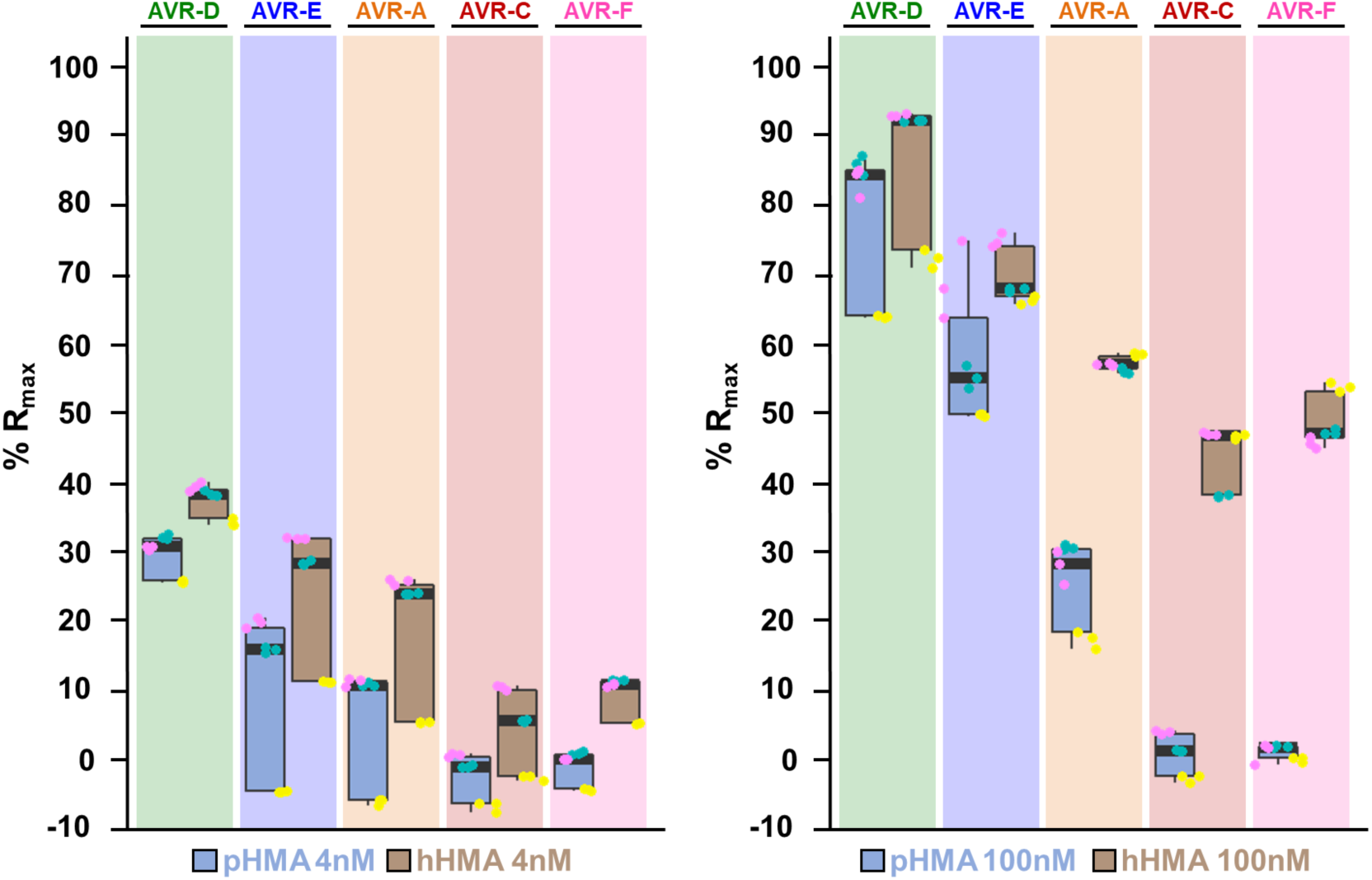
In vitro binding of Pikh-HMA domain to the AVR-Pik effectors measured by SPR is consistently higher compared to Pikp-HMA. Measurement of Pikp-HMA and Pikh-HMA binding to AVR-Pik variants measured by surface plasmon resonance. The binding is expressed as %R_max_ at HMA concentration of 4 nM (left) and 100 nM (right). Pikp-HMA and Pikh-HMA are represented by blue and brown boxes, respectively. For each experiment, three biological replicates with three internal repeats were performed and the data are presented as box plots. The centre line represents the median, the box limits are the upper and lower quartiles, the whiskers extend to the largest value within Q1 - 1.5× the interquartile range (IQR) and the smallest value within Q3 + 1.5× IQR. All the data points are represented as dots with distinct colours for each biological replicate.

**Supplemental figure 6.**
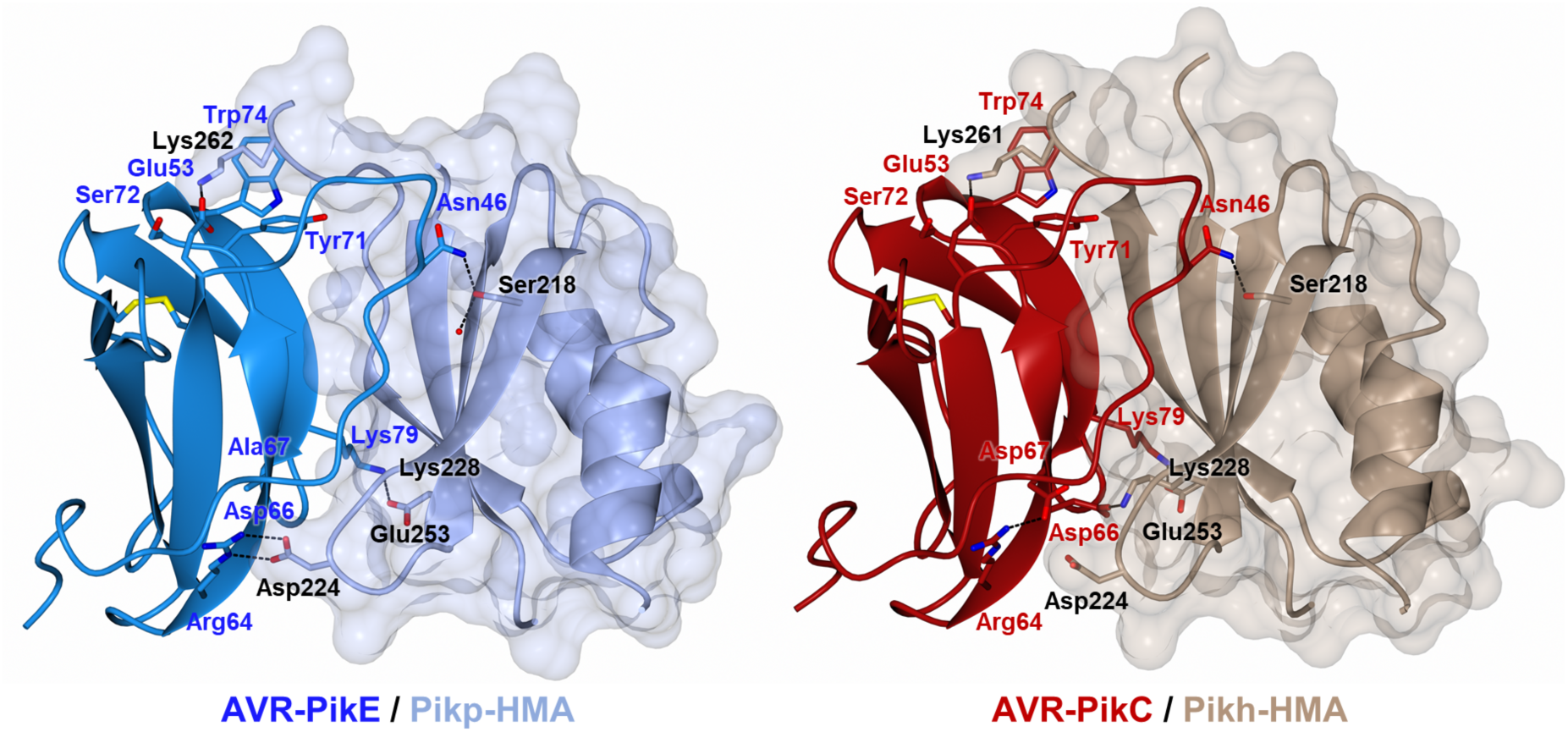
Overall structure of Pikh-HMA in complex with the AVR-PikC effector. Schematic representation of the structure of Pikh-HMA in complex with AVR-PikC (right). The structure of Pikp-HMA bound to AVR-PikE (PDB: 6G11) from [20] is included for comparison (left). HMA domains are presented as cartoon ribbons with selected side chains as cylinders; the molecular surface of the HMA domain is also shown. Pikh-HMA and Pikp-HMA are coloured in brown and ice blue, respectively. The effectors are shown in cartoon ribbon representation, with selected side chains as cylinders. AVR-PikC and AVR-PikE are coloured in crimson and bright blue, respectively. Hydrogen bonds/salt bridges are shown as black dashed lines and disulfide bonds as yellow cylinders. For clarity, of the two molecules of Pik-HMA present in the complex, only the one making extensive contacts with the effector is shown.

**Supplemental figure 7.**
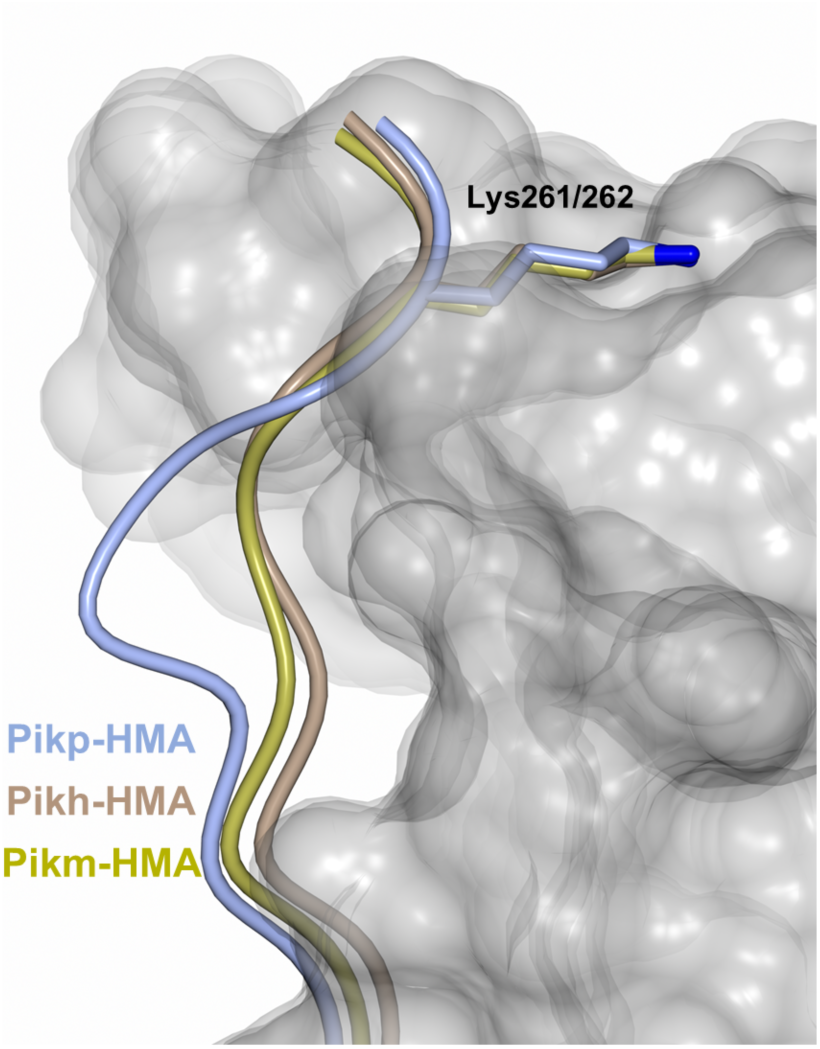
The Pikh-HMA domain adopts a conformation similar to Pikm-HMA. Superposition showing Pikp-HMA, Pikh-HMA and Pikm-HMA chains (coloured in blue, brown and yellow, respectively) bound to AVR-Pik. For clarity, only the Lys-261/262 side chain is shown. Two different effector alleles, AVR-PikE and AVR-PikC, are represented by their molecular surface coloured in grey.

**Supplemental figure 8.**
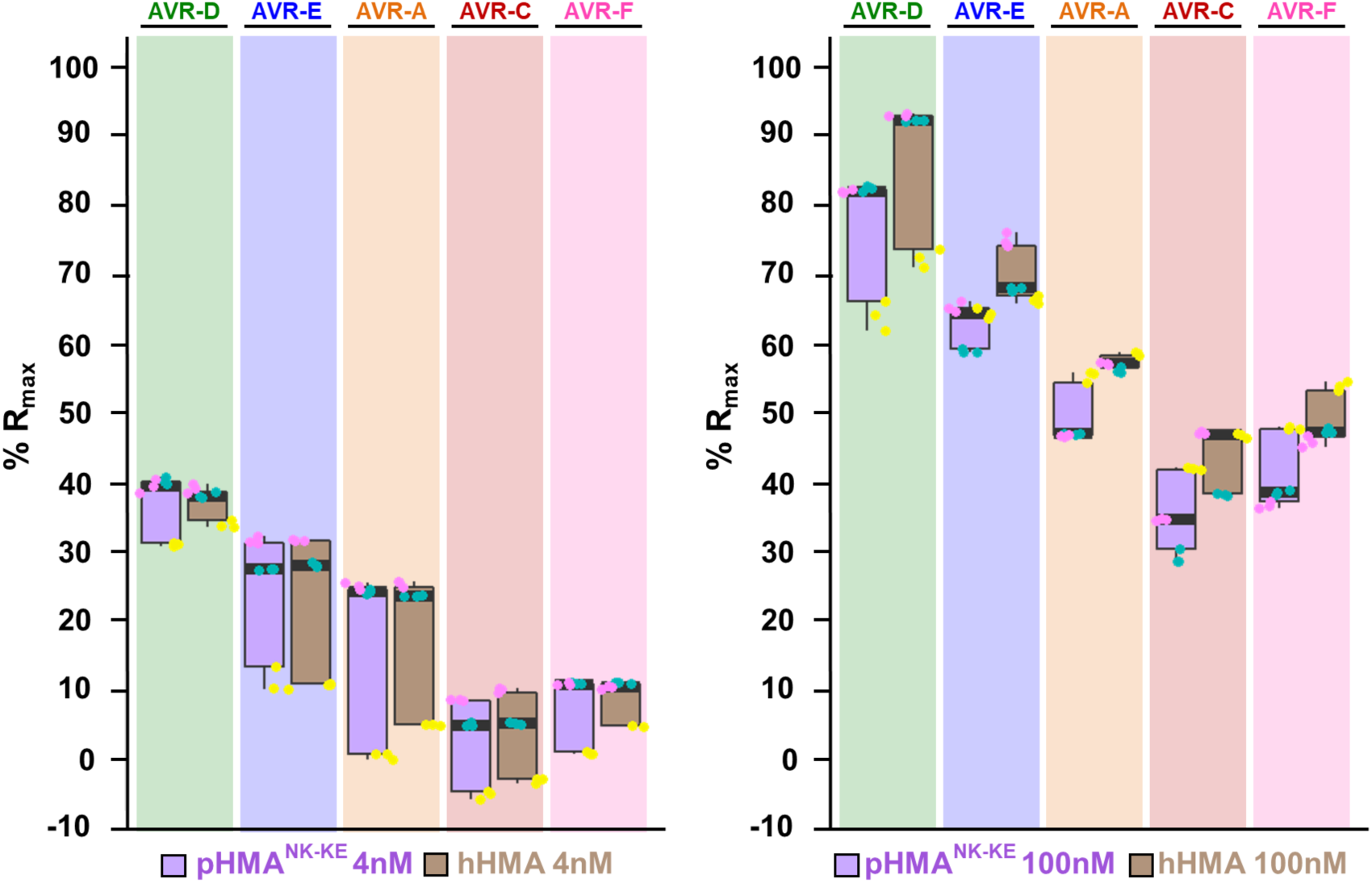
In vitro binding of the Pikh-HMA domain to the AVR-Pik effectors measured by SPR is similar to Pikp-HMA^NK-KE^. Measurement of Pikp-HMA^NK-KE^ and Pikh-HMA binding to AVR-Pik variants measured by surface plasmon resonance. The binding is expressed as %R_max_ at HMA concentration of 4 nM (left) and 100 nM (right). Pikp-HMA^NK-KE^ and Pikh-HMA are represented by purple and brown boxes, respectively. For each experiment, three biological replicates with three internal repeats were performed and the data are presented as box plots. The centre line represents the median, the box limits are the upper and lower quartiles, the whiskers extend to the largest value within Q1 - 1.5× the interquartile range (IQR) and the smallest value within Q3 + 1.5× IQR. All the data points are represented as dots with distinct colours for each biological replicate. Data for Pikh-HMA is also presented in **Supplemental figure 5** and were collected side-by-side at the same time.

**Supplemental figure 9.**
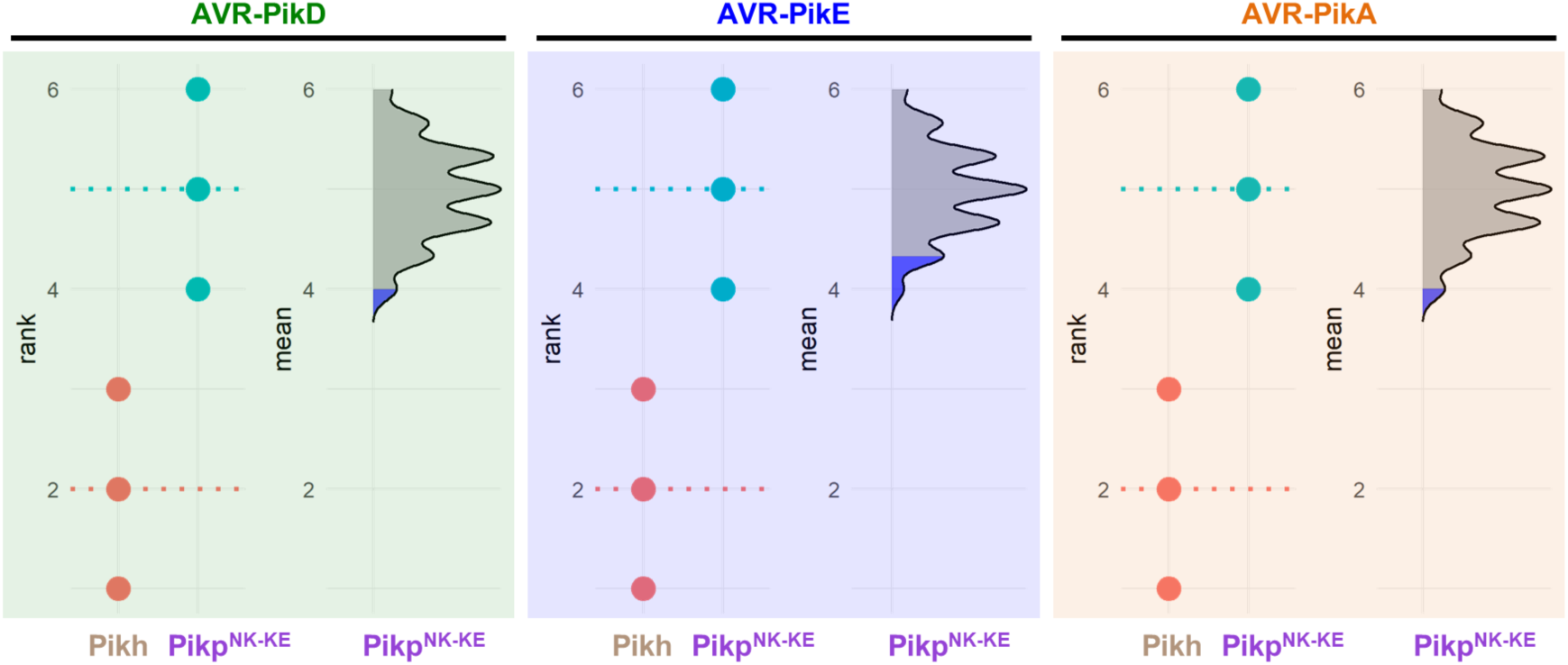
Estimation graphics for comparison of cell death mediated by Pikh and Pikp^NK-KE^. Statistical analysis by estimation methods of the cell-death assay for Pikh and Pikp^NK-KE^. For each effector, the panel on the left represents the ranked data (dots) for each NLR, and their corresponding mean (dotted line). The size of the dots is proportional to the number of observations with that specific value. The panel on the right shows the distribution of 1000 bootstrap sample rank means for Pikp^NK-KE^. The blue areas represent the 0.025 and 0.975 percentiles of the distribution. The responses of Pikh and Pikp^NK-KE^ are considered significantly different if the Pikh rank mean (dotted line, left panel) falls beyond the blue regions of the Pikp^NK-KE^ mean distribution.

**Supplemental figure 10.**
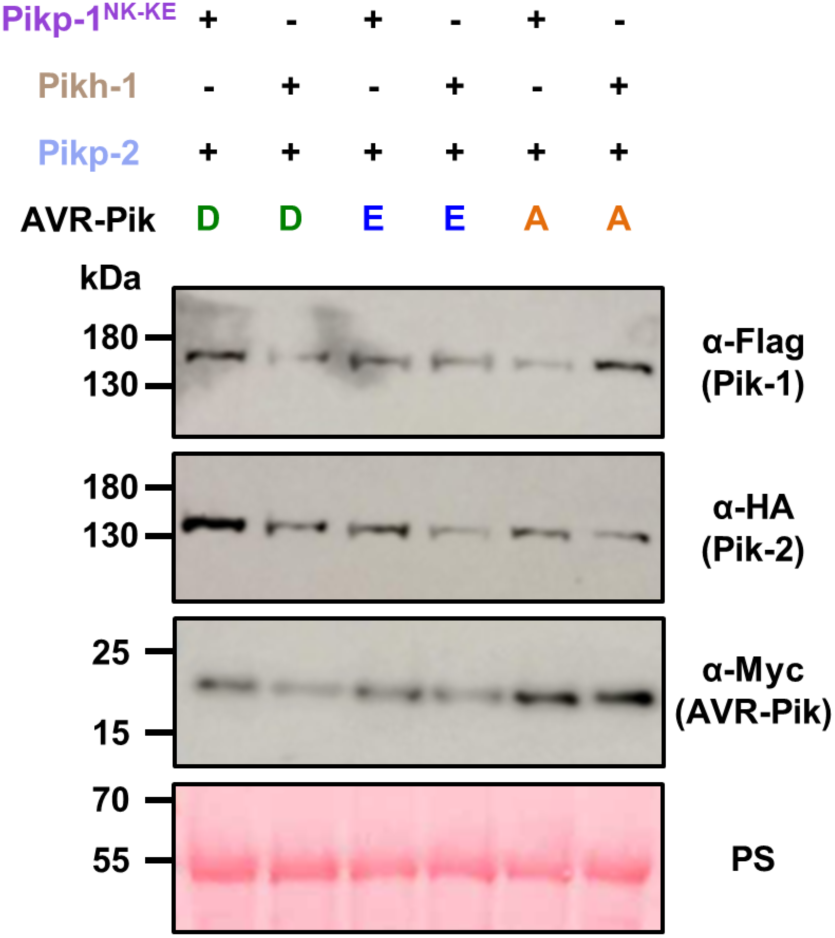
Western blots confirming the accumulation of proteins in *N. benthamiana*. Plant lysate was probed for the expression of Pikp^NK-KE^-1/Pikh-1, Pikp-2 and AVR-Pik effectors using anti-FLAG, anti-HA and anti-Myc antiserum, respectively. Total protein extracts were visualised by Ponceau Staining (PS).

**Supplemental table 1.**
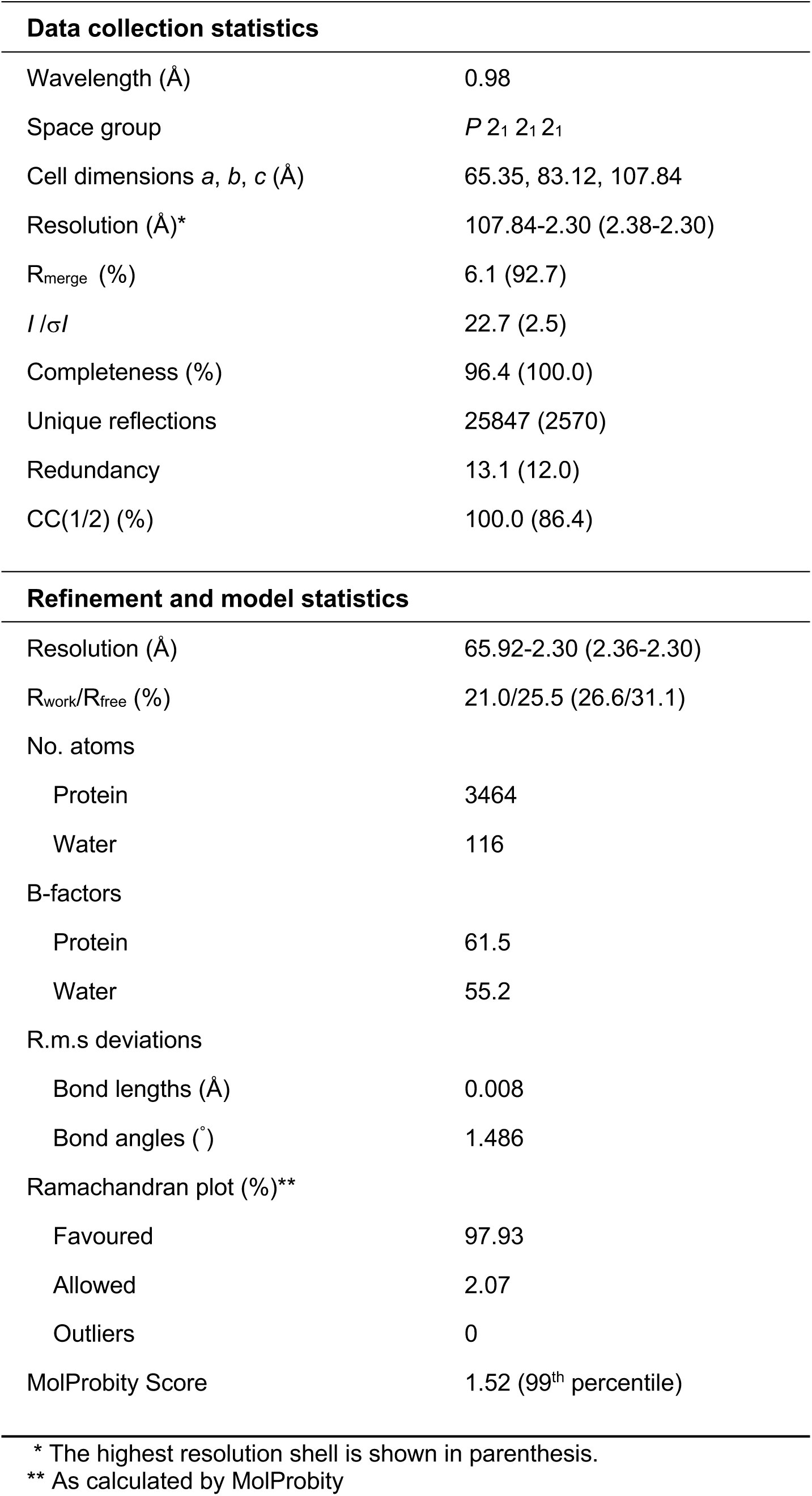
Data collection and refinement statistics for the crystal structure of the Pikh-HMA AVR-PikC complex.

**Supplemental table 2.** Summary of superposition analysis (as calculated with secondary structure matching (SSM) in CCP4MG version 2.10.10). For Pikh-HMA/AVR-PikC, chains E and F were used.

| Complex (compared<br>to Pikh/AVR-PikC) | AVR-Pik | HMA | Overall |
| --- | --- | --- | --- |
|  | r.m.s.d. - Å<br>(no. of residues) | r.m.s.d. - Å<br>(no. of residues) | r.m.s.d. - Å<br>(no. of residues) |
| Pikp/AVR-PikE | 0.31 (82 aa) | 0.61 (72 aa) | 0.70 (154 aa) |
| Pikm/AVR-PikE | 0.46 (81 aa) | 0.92 (74 aa) | 0.92 (165 aa) |

